# Paired single-cell multi-omics data integration with Mowgli

**DOI:** 10.1101/2023.02.02.526825

**Authors:** Geert-Jan Huizing, Ina Maria Deutschmann, Gabriel Peyré, Laura Cantini

## Abstract

The profiling of multiple molecular layers from the same set of cells has recently become possible. There is thus a growing need for multi-view learning methods able to jointly analyze these data. We here present Multi-Omics Wasserstein inteGrative anaLysIs (Mowgli), a novel method for the integration of paired multi-omics data with any type and number of omics. Of note, Mowgli combines integrative Nonnegative Matrix Factorization (NMF) and Optimal Transport (OT), enhancing at the same time the clustering performance and interpretability of integrative NMF. We apply Mowgli to multiple paired single-cell multi-omics data profiled with 10X Multiome, CITE-seq and TEA-seq. Our in depth benchmark demonstrates that Mowgli’s performance is competitive with the state-of-the-art in cell clustering and superior to the state-of-the-art once considering biological interpretability. Mowgli is implemented as a Python package seamlessly integrated within the scverse ecosystem and it is available at http://github.com/cantinilab/mowgli.

## Background

Single-cell sequencing technologies, providing a quantitative and unbiased characterization of cellular heterogeneity, are revolutionizing our understanding of the immune system, of development and of complex diseases^1–3^. A new frontier in the single-cell sequencing technologies is represented by multi-omics single-cell sequencing, allowing for the simultaneous profiling of multiple molecular readouts (e.g. transcriptome, chromatin accessibility, surface proteins) from the same cell^4–12^. Examples of these cutting-edge sequencing technologies are CITE-seq, simultaneously measuring RNA and surface protein abundance by leveraging oligonucleotide-conjugated antibodies^5^, and 10x Genomics Multiome platform, quantifying RNA and chromatin accessibility by microdroplet-based isolation of single nuclei.

Multi-omics single-cell sequencing platforms provide us with complementary molecular readouts from exactly the same set of cells, called in the following paired multi-omics data. The joint analysis of such data offers the exciting opportunity to understand how different molecular facets of a cell collaboratively define the cell’s function, morphology and state^13^. Several multi-view learning methods, jointly analyzing paired multi-omics data by taking into account their shared and complementary information, have thus been recently developed^14–23^. These methods, differently from unpaired integration ones^24,25^, take advantage of the known correspondences between cells across modalities. State-of-the-art multiview learning methods for single-cell multi-omics integration are based on integrative Matrix Factorization^14,19,22^, k-nearest neighbors^15^, or variational autoencoders^16–18^. Integrative Matrix Factorization (integrative MF) and variational autoencoders perform dimensionality reduction, jointly embedding the highdimensional multi-omics cellular profilings into a shared lower-dimensional latent space by leveraging common cells/observations^13,26^. Integrative MF, due to its linear nature, defines a latent space with a natural biological interpretation, but it is too simple to catch complex biological processes^13,26^. On the other hand, non-linear methods, as variational autoencoders, have shown great potential in clustering cells, but despite recent works on the subject^27,28^, they inherently lack biological interpretability. Improving integrative MF methods is thus crucial to striking a balance between interpretability and performance.

We here propose Multi-Omics Wasserstein inteGrative anaLysIs (Mowgli github.com/cantinilab/mowgli), a novel integrative MF method for single-cell multi-omics data combining integrative Nonnegative Matrix Factorization^29^ (integrative NMF) with Optimal Transport^30^ (OT). On one hand, Mowgli employs integrative NMF, popular in computational biology due to its intuitive *representation by parts* and further enhances its interpretability^29^. On the other hand, Mowgli enhances the clustering performances of integrative MF by taking advantage of OT, which we have previously shown to better capture similarities between single-cell omics profiles^31^.

We then extensively benchmark Mowgli with respect to the state-of-the-art in the integration of several paired multi-omics data profiled with CITE-seq^5^, 10X Genomics Multiome and TEA-seq^7^ platforms. Of note, while we focus on the integration of the currently available omics data, Mowgli can deal with paired multi-omics datasets with any type and number of omics, without any statistical assumption on the data. The performed in-depth comparison shows that Mowgli’s embedding and clustering quality outperform the state-of-the-art in controlled settings derived from real multi-omics data and are competitive with the state-of-the-art in more complex real multi-omics data. Of note, the latter are affected by the lack of an absolute groundtruth annotation on most real datasets. Finally, Mowgli is shown to improve the state-of-the-art in terms of biological interpretability through an in-depth biological analysis of TEA-seq data.

## Results

### Mowgli: a new tool for paired single-cell multi-omics data integration

We developed Multi-Omics Wasserstein inteGrative anaLysIs (Mowgli), a new tool for paired single-cell multi-omics data integration (github.com/cantinilab/mowgli).

Mowgli is based on integrative Matrix Factorization (integrative MF). Starting from *d* omics matrices 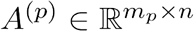 with *p* ∈ [1 … *d*], sharing the same columns (the cells) but having different features (e.g. genes, peaks), Mowgli jointly decomposes them into the product of omic-specific *dictionaries* 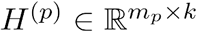 and a *shared embedding* 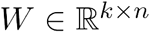 with *k* ≪ *m_p_* and *k* ≪ *n* (Figure 1A). As a standard nomenclature, in the following we will call *k* the number of *latent dimensions*, the columns of *H*^(*p*)^ *loadings* and the rows of *W factors*^26,32^.

**Figure 1.**
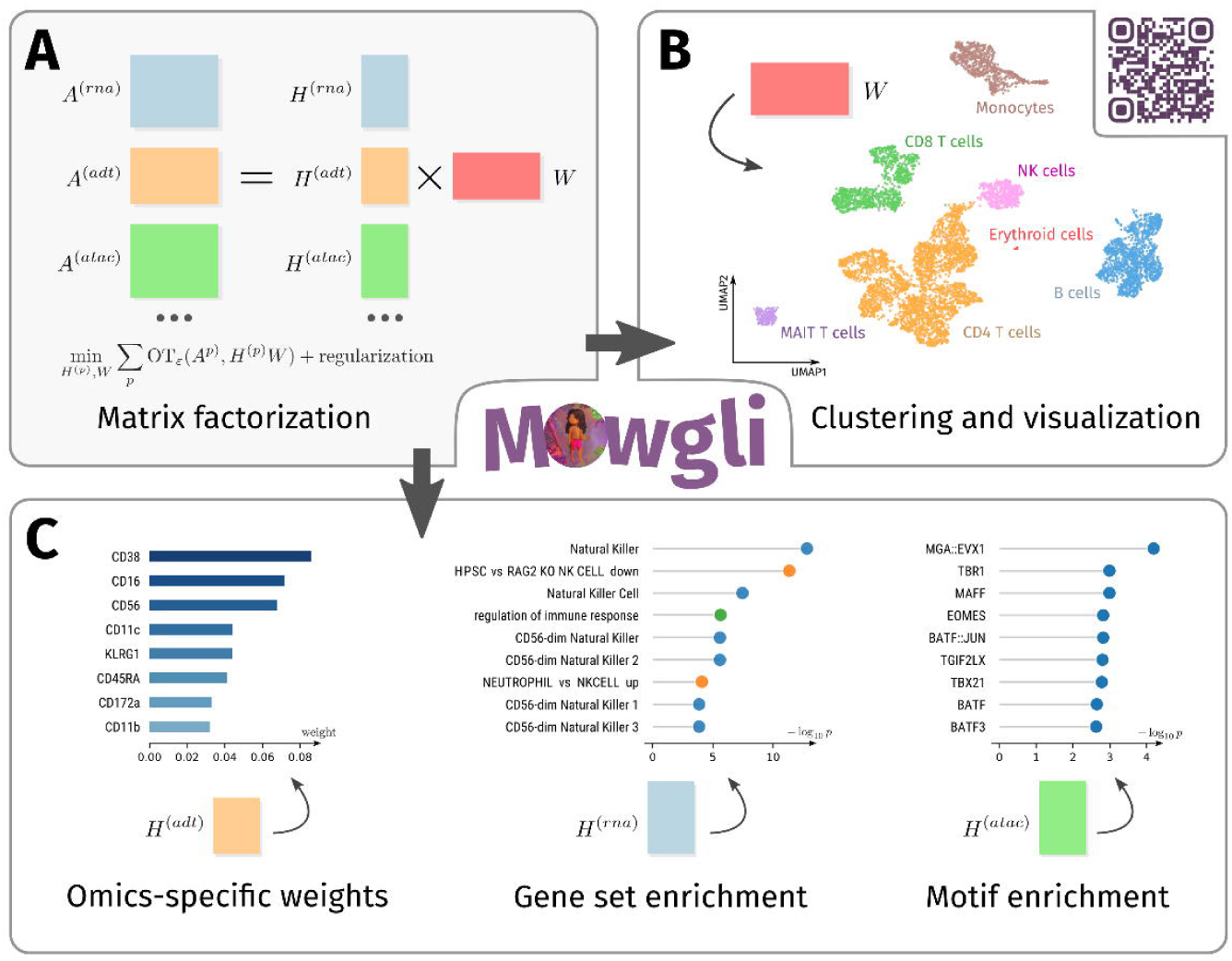
**(a)** Schematic visualization of Mowgli, an NMF-based model with an Optimal Transport loss; **(b)** the matrix *W* of Mowgli can be used for cell clustering and visualization; **(c)** The dictionaries *H*^(*p*)^ of Mowgli contain omics-specific weights for each latent dimension, which can be used for the biological characterization of the latent dimensions through gene set enrichment or motif enrichment analysis.

In line with state-of-the-art MF methods for multi-omics integration^33^, the cell embedding *W* can be used to visualize and cluster the cells (Figure 1B)^34–37^. The dictionaries *H*^(*p*)^ instead enable biological interpretation via gene set enrichment analysis^38^, motif enrichment analysis^39^, or by identifying markers among the top weights (Figure 1C).

The main innovation of Mowgli is to perform integrative MF by combining integrative Non Negative Matrix Factorization (integrative NMF) with Optimal Transport (OT). It thus solves the optimization problem:

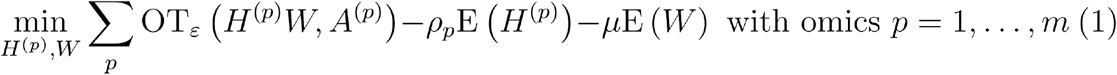

In computational biology, integrative NMF is usually applied with an Euclidean reconstruction term between *H*^(*p*)^*W* and *A*^(*p*) 19,22,24^. We here introduce instead a reconstruction term based on entropy-regularized Optimal Transport (OT) (see Eq1 and Methods), which unlike Euclidean or Kullback-Leibler losses leverages a notion of similarity between features. This choice is justified by the improved performance that we have previously observed once using OT to compare single-cell omics profiles^31^. Of note, outside of biology, OT has been already used in the reconstruction loss of NMF for the factorization of single matrices^40–42^ and single tensors^43^.

In addition, as in Rolet et al.^40^, we add to the optimization problem (Eq1) two entropic regularization terms *ρ_p_ E*(*H*^(*p*)^) and *μE*(*W*) (see Methods). These terms ensure that the loadings and embeddings are positive distributions and they control their sparsity (see Methods), a crucial feature to further enhance the known NMF’s “representation by parts” property^29^. *ρ_p_* and *μ* are the coldness parameters of *softmax* functions (see Methods) and thus offer a natural way to adjust sparsity. For instance, as *μ* approaches 0, cells will be assigned to only one factor. As instead *μ* increases, cells will be a combination of several factors. For all details on the mathematical formulation of Mowgli see Methods.

Of note, Mowgli is implemented as an open-source Python package seamlessly integrated into the classical Python single-cell analysis pipeline (github.com/cantinilab/mowgli). Users can thus take advantage of *scverse* tools like Scanpy and Muon for preprocessing and downstream analysis^44,45^. In addition, Mowgli provides a user-friendly visualization of top genes and enriched gene sets, thus helping biological interpretability.

In the following, we extensively benchmark Mowgli against the state-of-the-art: Seurat v4^15^ and MOFA+^14^. Although several methods exist^14–23^, we here focused on the leading methods for paired data integration that could be applied to the multiple combinations of single-cell omics data here considered. In addition, an integrative NMF baseline is also considered (see Methods), to further compare Mowgli with the standard integrative NMF.

### Mowgli’s cell embedding and clustering outperform the state-of-the-art in controlled settings derived from cell lines data

We first focused on evaluating Mowgli’s embedding and clustering performance in controlled settings derived from cell lines data. To represent a panel of realistic scenarios with different distributions of cells across three groups, we applied different transformations to a simple dataset composed of three cancer cell lines profiled with scCATseq (see Figure 2A). The scCATseq dataset provides a joint profiling of scRNA and scATAC from HCT116, HeLa-S3 and K562 cell lines^8^. Unlike simulated data, this solution allows us to avoid making assumptions on the distribution of the data. Indeed, generating simulated data following a Gaussian distribution, for instance, would favor methods that approximate single-cell data with this same distribution.

**Figure 2.**
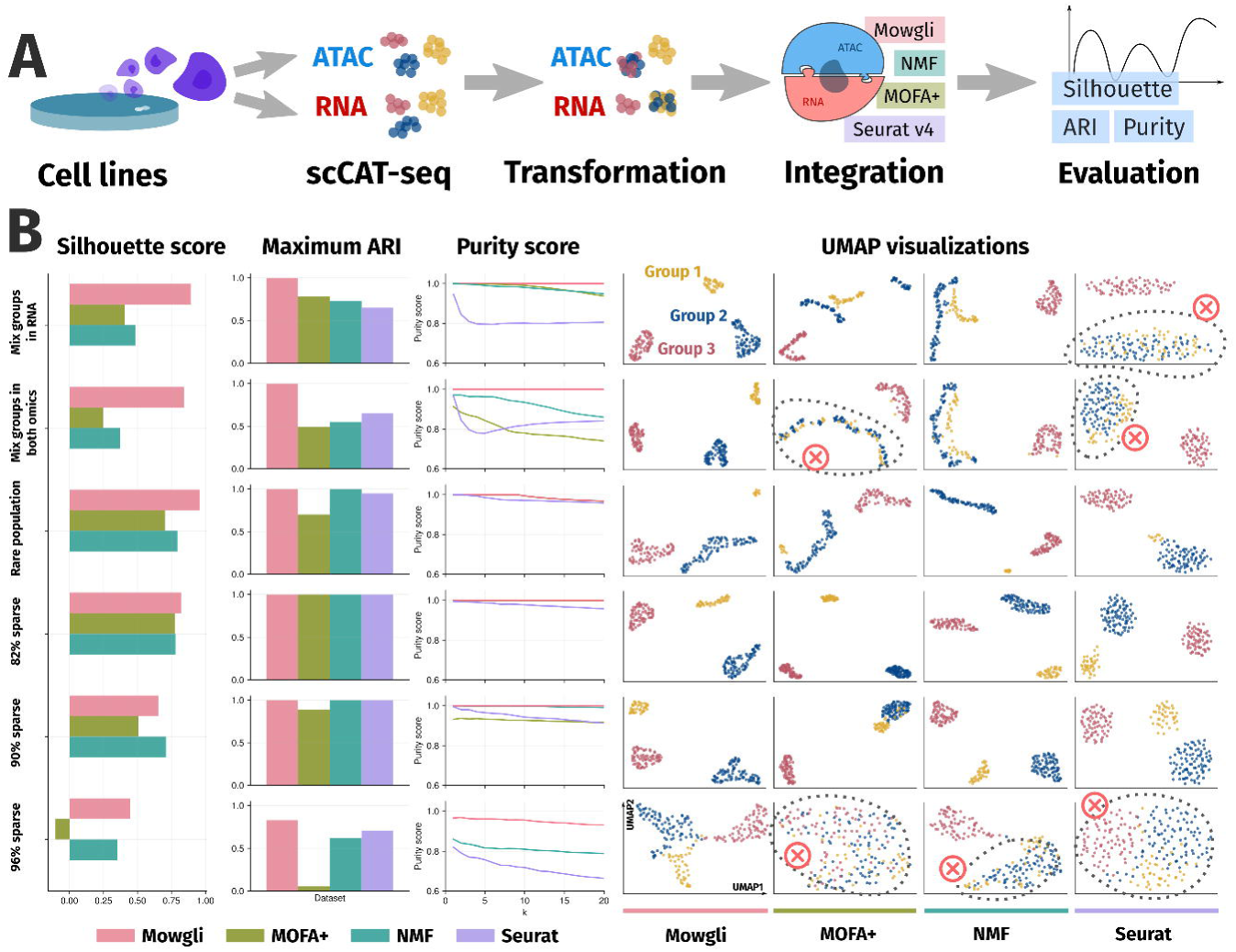
**(a)** Schematic representation of the benchmarking process; **(b)** The first three columns of this panel are devoted to silhouette scores, Adjusted Rand Indices (ARIs), and purity scores for the different methods on six controlled settings derived from cell lines data. The following four columns provide UMAP visualizations for the four benchmarked methods (Mowgli, MOFA+, NMF, Seurat v4) on six controlled settings derived from cell lines data. Different colors in these UMAP plots correspond to the three groups of cells imposed in the dataset.

The four scenarios in our panel represent distinct realistic challenges of multi-omics integration: (i) *Mixed in RNA* contains two cell populations that are mixed in scRNA, but well separated in scATAC; (ii) *Mixed in both* contains two cell populations mixed in scRNA and well separated in scATAC and two cell populations mixed in scATAC and well separated in scRNA; (iii) *Rare population* presents a population with much fewer cells than the others and (iv) *82% sparse, 90% sparse*, and *96% sparse* contain data with increasing percentages of dropouts (82-96%). Scenarios (i) and (ii) test the ability of methods to take into account the complementarity of different omic data. Scenario (iii) tests the ability of the methods to recover rare populations. Finally, scenario (iv) tests the robustness of the methods to dropout noise, while staying in a realistic range of dropouts for single-cell data. For details on the generation of these datasets see Methods.

We benchmarked Mowgli, Seurat v4, MOFA+ and integrative NMF based on natural metrics for embedding and clustering performance: silhouette score, Adjusted Rand Index (ARI), and purity score (see Methods). In addition, we computed UMAP visualizations for the different methods and datasets^35^.

As shown in Figure 2B, overall, Mowgli provides superior performance over the current state-of-the-art according to all metrics. Indeed, in all datasets except *90% sparse*, Mowgli has a performance greater or equal to that of other methods. In the *90% sparse* dataset, integrative NMF has a better silhouette score than Mowgli but the same ARI and purity score.

These performances are confirmed by looking qualitatively at the UMAP plots in Figure 2B. In *Mixed in RNA* and *Mixed in both* Seurat v4 confuses populations when individual omics are not sufficient to identify the three groups. Regarding dropouts, one of the most challenging features of single-cell data^46^, Mowgli shows the highest resilience with respect to the state-of-the-art. Indeed, while a sparsity of 96% is still coherent with realistic data^47^, MOFA+ and Seurat v4 confuse the three populations in the *96% sparse* dataset. On the opposite, Mowgli correctly separates the three groups of cells in *96% sparse*.

### Mowgli’s cell embeddings and clusterings are competitive with the state-of-the-art in complex and heterogeneous datasets

We then benchmarked Mowgli, Seurat v4, MOFA+ and integrative NMF based on their embedding and clustering performance on five paired single-cell multi-omics datasets (see Figure 3A). Of note, these data have been already largely used to benchmark single-cell multi-omics integrative methods^15,48,49^. The chosen datasets span different sequencing technologies, modalities, tissues and sizes: (i) *Liu* is a scCAT-seq cell lines dataset by Liu et al.^8^ (ii) *PBMC 10X* is a 10X Multiome human PBMC dataset from 10X Genomics (iii) *OP Multiome* is a 10X Multiome human bone marrow dataset from Open Problems^50^ (iv) *OP CITE* is a CITE-seq human bone marrow dataset from Open Problems^50^ (v) *BM CITE* is a CITE-seq human bone marrow dataset from Stuart et al.^25^. *BM CITE* is the larger dataset here considered, with 29,803 cells. Supplementary Table 1 lists the modalities, numbers of cells, and numbers of cell types for each dataset. For details on data preprocessing, see Methods.

**Figure 3.**
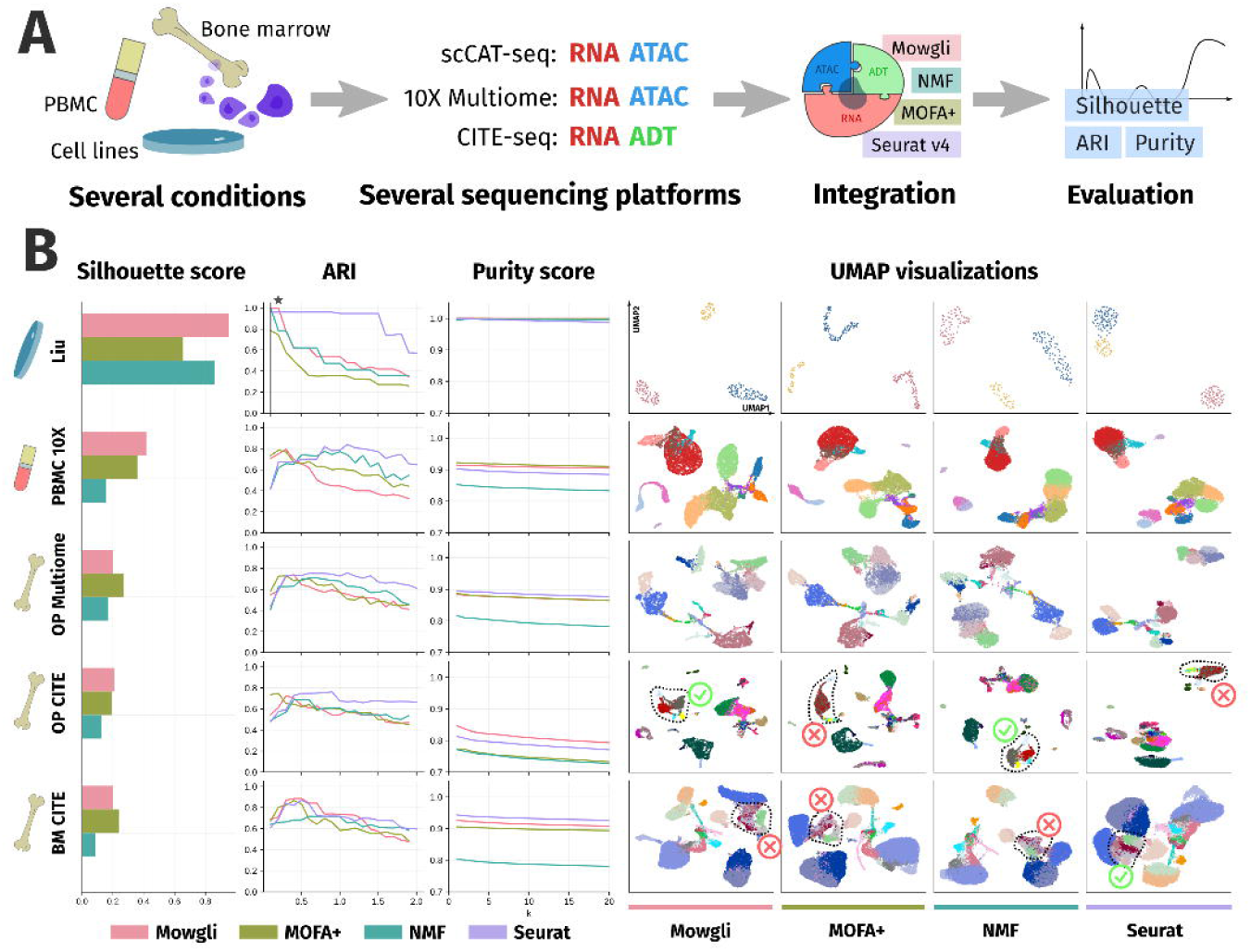
**(a)** Schematic representation of the benchmarking process; **(b)** The first three columns of this panel are devoted to silhouette scores, Adjusted Rand Indices (ARIs), and purity scores for the different methods on five complex paired single-cell multi-omics data already largely used to benchmark integrative methods. The following four columns provide UMAP visualizations for the four benchmarked methods (Mowgli, MOFA+, NMF, Seurat v4) on the same data. Different colors in these UMAP plots correspond to the different ground-truth cell type annotations provided with the data.

We benchmarked Mowgli, Seurat v4, MOFA+ and integrative NMF based on the same natural metrics used in the previous section. Since these metrics require a groundtruth annotation, we used the cell-type annotations available from the original publications of these data. In *Liu* the ground-truth annotations are based on the cell line of origin and thus well-defined. On the contrary, the annotations of the other datasets were computationally derived, thus biasing the evaluation toward methods closest to their annotation pipeline. For instance, the *BM CITE* annotation is obtained by projecting the dataset onto an atlas using Seurat v3^25^.

As displayed in Figure 3B, Mowgli can handle large single-cell datasets and deliver embedding and clustering performances competitive with the state-of-the-art, especially considering the lack of absolute ground-truth annotations on most datasets here employed.

In particular, according to the silhouette score, Mowgli outperforms other methods in *Liu, PBMC 10X*, and *OP CITE*. MOFA+ performs best in the other two datasets. In terms of ARI score across resolutions, Seurat v4 performs best in *PBMC 10X, OP Multiome*, and *OP CITE*. Mowgli and Seurat v4 perform comparably in the *BM CITE* dataset. In the *Liu* dataset, only MOFA+ and Mowgli reach a maximum ARI of 1. Of note, in *Liu*, ARIs should be compared only at low resolution, as higher resolutions lead to overclustering. In terms of purity score, Mowgli outperforms other methods in the *OP CITE* dataset, and it is comparable to MOFA+ in the *PBMC 10X* dataset. Finally, the purity scores of all methods are comparable in the *Liu* dataset.

The UMAP plots in Figure 3B give a qualitative intuition of the described performance. In *OP CITE*, only integrative NMF and Mowgli correctly separate subpopulations of B cells (Figure 3B circled). In *BM CITE*, MAIT T-cells and subpopulations of CD8+ T-cells (Figure 3B circled) are more neatly separated in Seurat v4 than in other methods. However, as explained previously, the annotation pipeline of *BM CITE* might favor Seurat v4.

### Mowgli improves the biological interpretability of the state-of-the-art by providing cell-type specific factors in TEA-seq data

We benchmarked Mowgli with respect to MOFA+ based on its biological interpretability (see Figure 4A). Indeed, MOFA+ is the leading single-cell multi-omics integration tool providing a user-friendly biological interpretability of its latent dimensions^14^.

**Figure 4.**
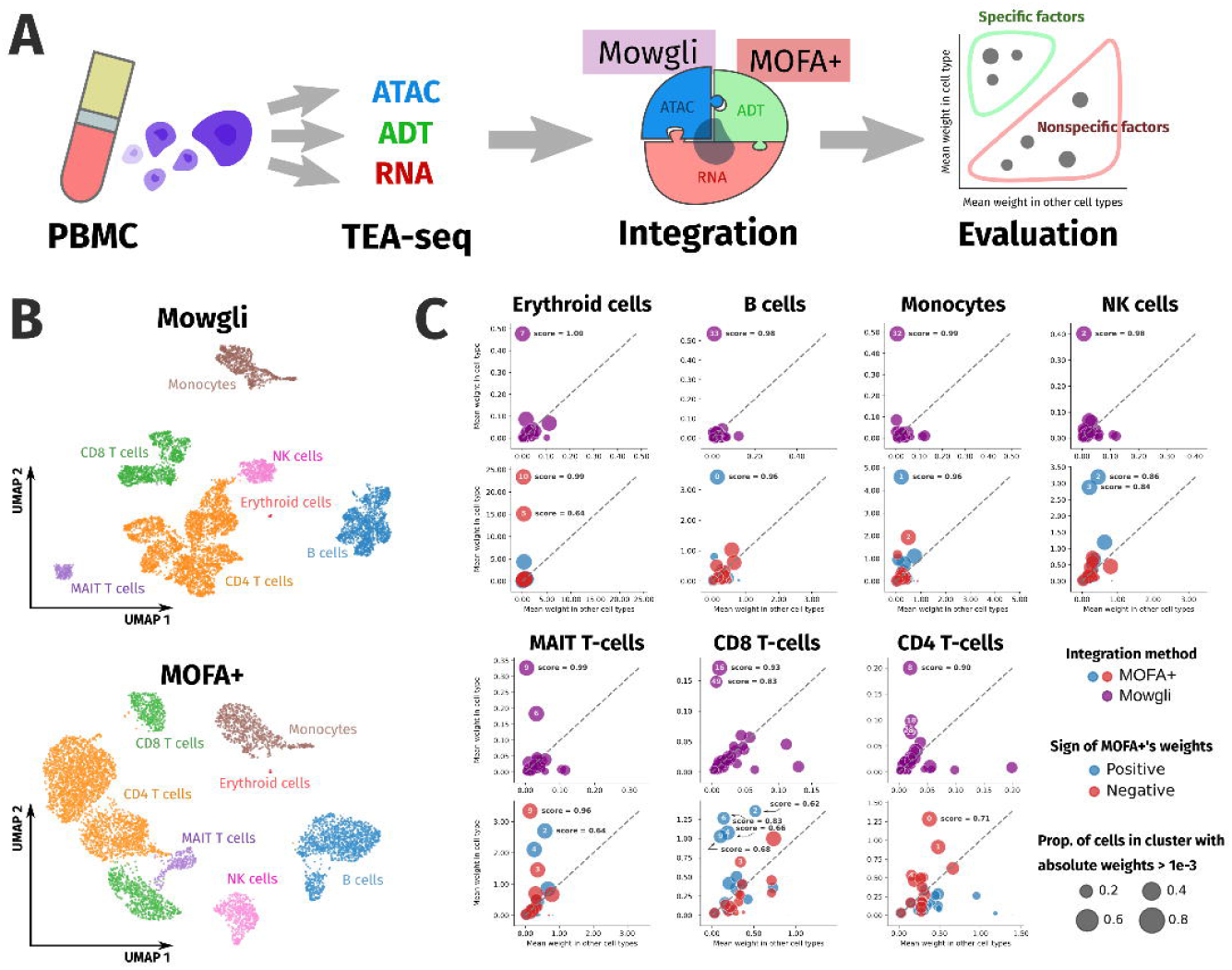
**(a)** Schematic representation of the evaluation process on biological interpretability; **(b)** UMAP visualization of Mowgli’s and MOFA+’s embeddings. The colors correspond to a marker-based cell-type annotation of the cells; **(c)** average weights within and outside of a cell type are plotted for each factor of Mowgli (violet) and MOFA+ (red for the negative part and blue for the positive one). For each cell type, the best specificity scores are reported in bold.

For this benchmark we considered a TEA-seq dataset of human PBMCs, corresponding to the paired profiling of: scRNA-seq, scATAC-seq, and surface proteins^7^. This dataset allows us to test the methods on more than two omic datasets, thus taking into account more complementary layers of molecular regulation.

First, MOFA+ and Mowgli were independently applied for the integration of the three omics constituting the TEA-seq data. As the dataset was not provided with an annotation of the cells, we separately clustered the embeddings obtained from Mowgli and MOFA+’s and annotated them based on gene and protein markers (see Supp Figure 1, see Figure 4B). We identified in this way coarse immune cell types: CD4 T-cells, CD8 T-cells, B cells, Natural Killer (NK) cells, MAIT T-cells, Monocytes and Erythroid cells. Of note, the cell type annotations obtained with the two tools agree at 97%, and match an independent RNA-based annotation obtained through Azimuth (see Supp Figure 2). Both methods are thus able to recover the expected cell-types through clustering of their embeddings.

To then test the biological interpretability of Mowgli and MOFA+, we evaluated the specificity of the associations between their factors and the identified immune cell types. The underlying assumption we are making here is that an interpretable method should provide factors that are not broadly active in all the cells, but selectively associated to a cell type. Indeed, characterizing a cell type which results from a combination of many factors is a daunting task. On the contrary, having cell type-specific factors makes the biological characterization of the associated cell type straightforward. To evaluate such specificity, for each cell type, we plotted how the Mowgli and MOFA+ factors are distributed according to their mean weight within the cell type and their mean weight outside the cell type (Figure 4C). Factors specific to a cell type should have a high average weight within the cell type and a low average weight outside the cell type, thus falling in the upper left corner of the plots. As MOFA+’s factors are not constrained to be positive and their positive and negative parts could be associated with different biological information, we split each factor into two parts, as done in MOFA+’s interpretation tools^14^. In addition, we quantified the performance of each factor with a specificity score, also reported in bold in Figure 4C, and defined in the Methods section.

As shown in Figure 4C, while MOFA+ tends to associate multiple factors to the same cell type, Mowgli frequently defines clear one-to-one associations between factors and cell types. In addition, the specificity score of such factors is higher in Mowgli than in MOFA+. This is particularly striking in NK cells, MAIT T-cells, CD8 T-cells and CD4 T-cells, where MOFA+ seems to aggregate information from many factors whereas Mowgli is more selective. Of note, as shown in Supp Figure 3, the multiple factors associated by MOFA+ to the same cell type do not necessarily correspond to subpopulations of the same cell type.

### Mowgli identifies relevant subpopulations of immune cells in TEA-seq data

We finally focused on the biological relevance of the factors identified by Mowgli on the human PBMC TEA-seq data, described in the previous section. Indeed, while in the previous section we only considered coarse immune cell types (e.g. B cells, CD4 T-cells, CD8 T-cells), Mowgli could identify multiple factors able to subset such cell types into relevant subpopulations (Figure 5 A,B; Supp Figure 4). For example, Mowgli identifies factors splitting the B cell cluster into two subpopulations: memory and naive B cells. In the same way, Mowgli detects factors associated to CD8 T-cells subpopulations (naive, central memory and effector memory), monocytes subclusters (classical and non-classical), dendritic cells subpopulations (plasmacytoid and conventional) and Natural Killer (NK) subclusters (CD56^dim^ and CD56^bright^). The association of the factors with specific immune subpopulations is here made based on top ranked genes and proteins in effector memory CD8 T cells, naive B cells, memory B cells and CD56^dim^ NK cells. For all other populations the association with factors is instead based on the correlation of the factors’ weights with that of known protein markers. Figure 5B displays side-by-side the UMAP plots showing the similarity between the distribution of the factors’ weights and the activity of the protein markers of their associated immune subpopulations. The UMAP visualizations of all marker proteins and all factors are available in Supp Figure 5.

**Figure 5.**
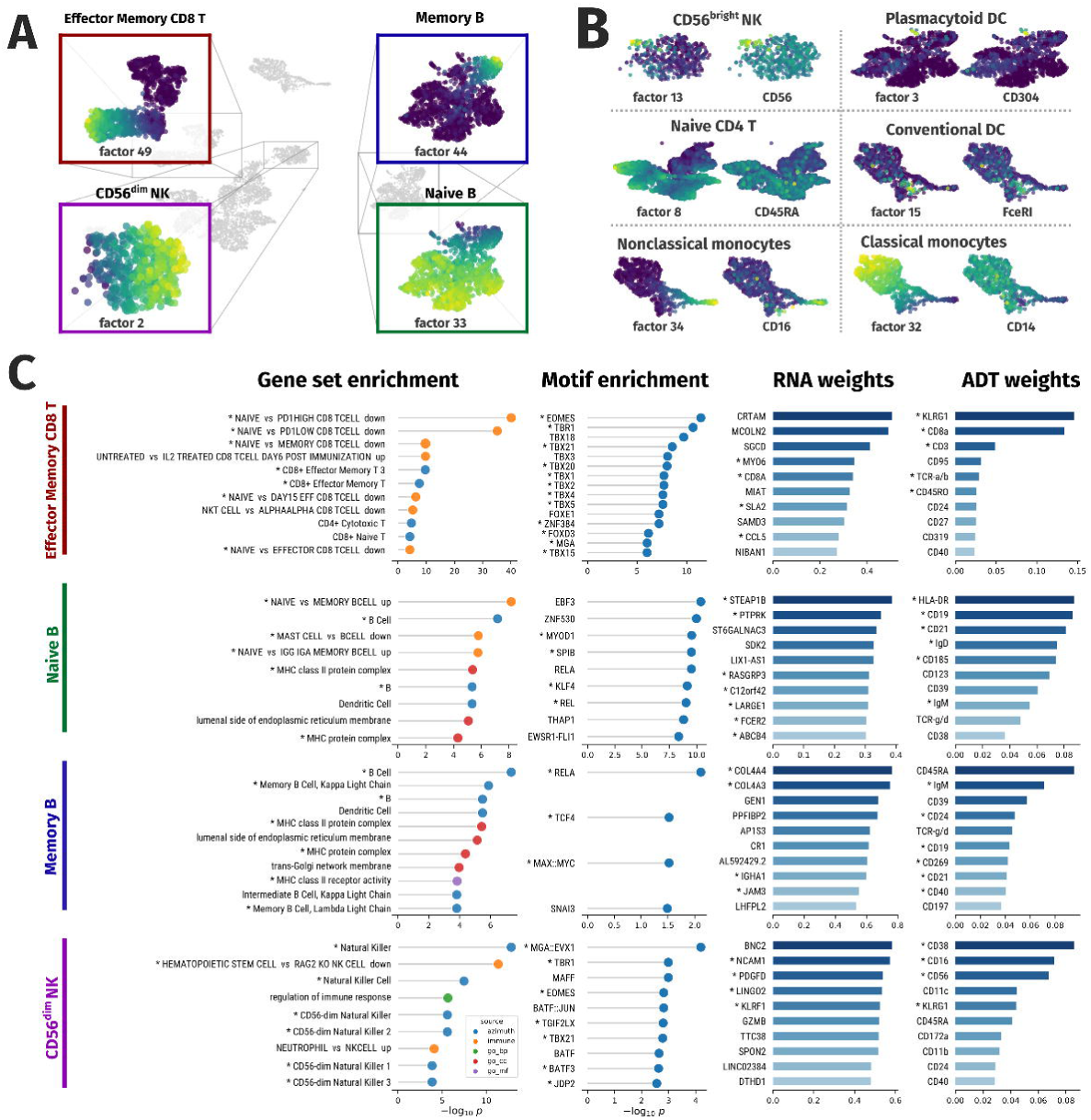
**(a)** UMAP visualization of Mowgli’s embedding with focus on four specific immune subpopulations (Effector Memory CD8 T-cells, memory B cells, CD56^dim^ NK cells, naive B cells) for which the UMAP is colored based on factor weights; **(b)** UMAP visualization of Mowgli’s embedding colored by factor weight and protein marker weight for other factors corresponding to specific subpopulations of cells; **(c)** Top genes, proteins, gene sets and Transcription Factors (TFs) for the 4 factors visualized in panel a. Stars denote gene sets and markers pertinent for the immune subpopulation associated with the factor and TFs targeting the top genes.

These same results could not be obtained with MOFA+, due to its lower biological interpretability observed in the previous section. In MOFA+, factors having similar patterns of those observed in Mowgli could be obtained for effector memory CD8 T-cells, memory B cells, non-classical monocytes and CD56^dim^ NK cells (see Supp Figure 3 and Supp Figure 6). For all other immune subpopulations identified by Mowgli, no factor having a similar pattern could be obtained in MOFA+. As a consequence, interpreting with MOFA+ the pathways associated to CD56^bright^ NK cells, for example, would require to complexly combine the pathway enrichments obtained from different factors. On the contrary, the same analysis in Mowgli can be easily realized by looking at the pathways enriched in the loadings of its 13th factor.

Finally, we looked at the biological information that Mowgli could provide regarding the identified immune subpopulations. For this part, we focused on the factors associated with four immune cell subpopulations: effector memory CD8 T-cells (factor 49), naive B cells (factor 33), memory B cells (factor 44), and CD56^dim^ NK cells (factor 2). For each of these four factors, we considered their associated loadings in *H*^(*rna*)^, *H*^(*adt*)^ and *H*^(*atac*)^ and analyzed the top genes in *H*^(*rna*)^, top proteins in *H*^(*adt*)^, the gene sets enriched in *H*^(*rna*)^ and the motifs enriched in *H*^(*atac*)^ to verify the biological information that could be extracted from the output of Mowgli (see Methods). Figure 5C displays the results obtained from this analysis.

For effector memory CD8 T-cells (CD8 TEM cells), corresponding to factor 49, Mowgli could extract two top genes (CRTAM and KLRK1), known to be essential for CD8+ T-cell-mediated cytotoxicity^51,52^, two top proteins (CD45RO, TCR-a/b) that are a known memory T cell marker and a T cell receptor, respectively^53,54^. More interestingly, also several Transcription Factors (TFs) candidate regulators of this subpopulation are identified, among them EOMES and TBX21 (aka T-bet), known to be important for CD8 TEM development^55^. In addition, five of the top candidate TF regulators (TBR1, TBX21, TBX4, TBX5 and MGA) target three of the top genes of the same factor (CCL5, CRTAM, and IL21R), thus suggesting a regulatory program possibly important for CD8 TEM cells.

In naive B cells (factor 33), Mowgli identifies as top genes FCER2 (aka CD23), a low-affinity receptor for immunoglobulin E (IgE) with an essential role in the differentiation of B-cells^56^ and MARCH1, which downregulates the surface expression of major histocompatibility complex (MHC) class II molecules^57^. In the top proteins we can single out CD19, CD21, and HLA-DR, well-known markers of B cells^58^. In addition, the relative weights of IgD and IgM in factor 33 are coherent with the repartition already described for naive B-cells^58^. Finally, among the top TF candidate regulators of factor 33, Early B-cell Factors (EBF3 and EBF1)^59^ and NF-kB proteins (REL and RELA) stand out as regulators of the top genes of the same factor. Of note these TFs play an essential role in B-cell development, maintenance, and function^60^.

For memory B cells (factor 44), Mowgli extracts as top genes: IGHA1 and IGKC, part of immunoglobulin complexes^61^ and JAM3, belonging to the Immunoglobulin superfamily and already studied in the context of B cell homing and development^62–64^. The top proteins include the well-known B cell markers CD19, CD21, and HLA-DR^58^. In addition, as observed before for naive B cells, the relative weights of IgD and IgM in factor 44 are coherent with the repartition already described for memory B-cells^58^. In the top TFs emerging from our motif analysis and targeting the top genes we finally find RELA, TCF4 and MAX::MYC, known to be involved in the transcriptional regulation of memory B cell differentiation^65^.

Finally, in CD56^dim^ NK cells (factor 2), Mowgli detects at top genes: NCAM1 (aka CD56), the go-to marker for NK cells^66^; KLRF1 and KLRD1, genes of the KLR family of receptors controlling NK cell activity^67^; GZMB, involved in NK-cell mediated cytotoxicity^68^; SLAMF7, mediating NK cell activation^69^. Top proteins include CD56, the canonical marker of NK cells^66^, but its weight is lower than that of CD16, which is coherent with the expression profile of CD16+CD56^dim^ NK cells^66^. Regarding TF candidate regulators, we detect EOMES and TBX21 (aka T-bet), which are critical to NK-cell differentiation^70^, Maf-F, having a key role in the regulation of NK cell effector functions by IL-27, and JUNB::FOSB, early activator protein (AP)-1 TFs that regulate NK-meditated cytotoxicity^71,72^. Finally, a strong regulatory program seems to emerge here with four of the top candidate TF regulators for factor 2 (MGA::EVX1, EOMES, TBX21, and JDP2) targeting four of the top genes of the same factor (C1orf21, IL18RAP, PTGDR and SLAMF7).

## Conclusions

Multiple technologies allowing the multi-omics profiling of the same set of cells are currently available. We thus need integration methods able to jointly learn from multiple omics data profiled on the same cells.

In this article we introduced Multi-Omics Wasserstein inteGrative anaLysIs (Mowgli), an integrative method for paired multi-omics data that enables rich biological interpretation for any type and number of omics. We then in-depth benchmark Mowgli’s cell embedding and clustering performance with respect to the state-of-the-art in controlled settings derived from scCAT-seq profiling of cancer cell lines. Mowgli outperforms in this benchmark the state-of-the-art showing its high potential even in challenging conditions. We then considered more complex and heterogeneous data profiled with CITE-seq and 10x Genomics Multiome technologies. On these data, Mowgli performed comparably with the state-of-the-art, with no method clearly outperforming others. Finally, regarding the biological interpretability, once tested on TEA-seq data, corresponding to paired scRNA, scATAC, and surface protein profiling, Mowgli produces biologically meaningful representations superior to those of the state-of-the-art.

A major limitation affecting this benchmark and all other focused on paired multi-omics integration corresponds to the lack of a high-quality biological annotation of the cells. While in some cases Fluorescence-activated cell sorting (FACS) could represent a clear solution for an independent annotation of the cells, paired multi-omics data with this type of annotation are lacking in the literature.

Concerning then Mowgli’s limitations and possible future extensions, it would be interesting to extend Mowgli to deal with batch correction once integrating paired multi-omics data. Indeed, most recent large scale paired multi-omics data are profiled in different centers thus creating batch correction issues. In addition, Mowgli does not contain a straightforward approach to define the number of latent dimensions. This problem has been however extensively studied in NMF literature and users can rely on classical tools like the cophenetic coefficient or the elbow method. At the same time, Mowgli is fairly robust to changes in the number of latent dimensions (*k*) thus suggesting that small changes in *k* will not affect its performance. Finally, as OT is inherently expensive to compute, Mowgli requires GPU computations for the larger datasets presented in this article. However, the availability of GPUs is nowadays a standard in research centers and this will be further enhanced in the future once larger single-cell datasets will be available.

## Methods

### Notations

Let us consider *n* cells, measured across several modalities. Each modality *p* has *m_p_* features (e.g. genes). Let us denote 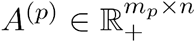 the dataset for modality *p*. Additionally, we impose each column of *A*^(*p*)^ to sum to 1, i.e. be a discrete probability distribution.

### Optimal Transport

Optimal Transport (OT), as defined by Monge^30^ and Kantorovich^73^, aims at finding a coupling *P* between two probability distributions *a* and *b* that minimizes the cost of transporting one distribution to the other. In the discrete case, the classical OT distance, also known as the Wasserstein distance, between *m*-dimensional histograms *a* = (*a*_1_, …, *a_m_*) and *b*, …, *b_m_*) is defined as

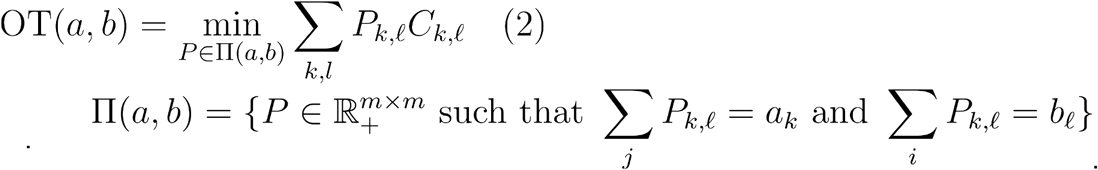

where

The coupling *P* ∈ Π(*a, b*) represents how the mass in the discrete probability distribution *a* is moved from one bin (e.g. gene) to another one in order to transform *a* into *b*.

The ground cost 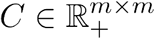 is a pairwise distance matrix that encodes the penalty for transporting mass from one feature (e.g. gene) to another. Hence, *C* should be chosen in such a way that similar bins (e.g. genes) *k* and *ℓ*, have a low cost *C_k,ℓ_*. Here, for a certain omic *p*, we define *C* in a data-driven way as the matrix of pairwise Pearson correlation distances between the features, i.e. the rows in our dataset *A*^(*p*)^. In other words, denoting 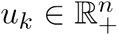 and 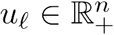 two rows in our dataset,

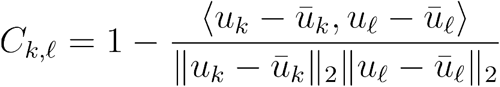

where 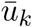 is the mean of the elements of *u_k_* and 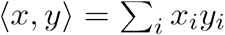 is the dot product. This choice of ground cost gave best results in our previous work^31^.

Due to the high-dimensionality of single-cell data, we use the entropic regularization of OT, a fast and GPU-enabled approximation of classical OT computed using the Sinkhorn algorithm^74^:

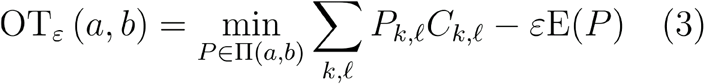

where the entropy *E* is defined as 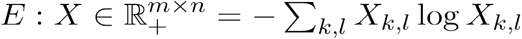.

If *ε* is set to zero, (Eq3) corresponds exactly to classical OT (Eq2). Increasing values of *ε* correspond to a more diffused coupling *P*. In previous work, we showed the entropic regularization of OT to improve similarity inference between single-cell omics profiles compared to classical notions of distance^31^.

As explored in^31^, entropic regularization is expected to control the systematic noise due to technical dropouts and to the stochasticity of gene expression at the singlecell level. In addition, more diffused couplings increase the exchange of mass between features. This enables OT to leverage the relationships between features (e.g. genes), motivating further its application to single-cell data.

### Mowgli

We aim to decompose each matrix *A*^(*p*)^ as the product of a matrix 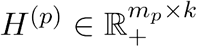 (the modality-specific dictionaries) and 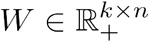 (the embeddings, shared across modalities). The integer *k* is the number of dimensions of the latent space and should be small compared to the number of features. We use the entropic regularization of OT as a reconstruction loss to compare *H*^(*p*)^*W* to the reference data *A*^(*p*)^.

In addition, we require the columns of *H*^(*p*)^*W* to sum to 1, i.e. belong to the simplex. We thus impose that the columns of *H*^(*p*)^ sum to one, and that the columns of *W* sum to one. Following Rolet et al.^40^, we use the entropy function E defined previously, with a value of –∞ when columns do not sum to one.

Combining the reconstruction and the entropy terms yields the loss

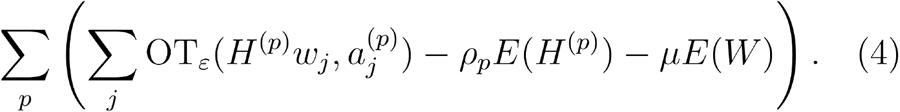

Note that for the sake of readability, we write OT_*ε*_ for all *p*, but this loss actually depends on an omic-specific ground cost *C*^(*p*)^, which itself depends on *A*^(*p*)^ (see Eq3). The parameters *ρ_p_, μ* control the sparsity of the columns of *H*^(*p*)^ and *W*. In order to make these parameters more comparable across omics and datasets, we define 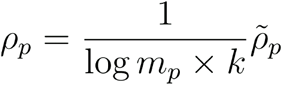 and 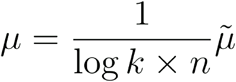.

Default values of 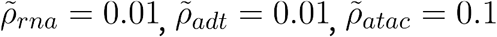, and 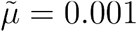 yielded best results across experiments. For *Liu* and datasets derived from *Liu*, we run the method with 5 factors. For other datasets, we choose 50 factors (see Supp Figure 7).

Similarly to Rolet et al.^40^, we alternate between minimizing (Eq4) on *H*^(*p*)^ and *W*. One can show that these smooth minimization problems on *H*^(*p*)^ and *W* are equivalent to the following smooth minimization problems on new *dual* variables *G*^(*p*)^. These problems can be solved using standard optimization methods, and the method of choice is L-BFGS, a limited-memory quasi-Newton method.

- *Optimizing H*^(*p*)^. We solve the following smooth minimization problem:

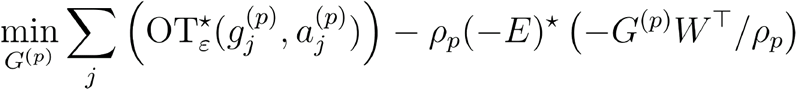 Then, we update the primal variable as follows:

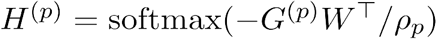 The column-wise softmax is defined as:

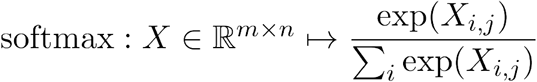
- *Optimizing W*. We solve the following smooth minimization problem:

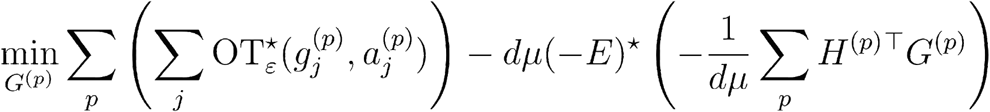 Then, we update the primal variable as follows:

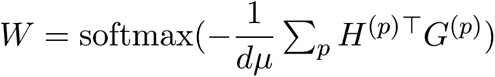

Here, OT* and (−*E*)* denote the Legendre duals of the OT_*ε*_ and –*E* functions, and their smooth closed form expressions are defined in Rolet et al.^40^. In the subproblems defined above, the coefficients *ρ_p_* and *μ* parametrize *softmax* functions, and hence control the sparsity of distributions. When *ρ_p_* tends to zero, the *softmax* behaves like an *argmax* and the distributions tend to Diracs.

The code is implemented in Python and relies on PyTorch^75^ for matrix operations on the GPU and on Muon^45^ and Scanpy^44^ to handle single-cell multimodal data.

### MOFA+

We compare Mowgli to MOFA+^14^, a variational inference method analogous to sparse PCA for multi-omic data. We use the R interface MOFA2 with default training parameters. MOFA+ provides a parameter drop_factor_threshold designed to keep only informative factors, but we found that in practice it removed important information. For example, the benchmark in Zuo and Chen^18^ only kept one factor for MOFA+, which is not enough to represent cellular heterogeneity in the data. We thus choose to keep 5 factors for *Liu* and the datasets derived from *Liu*, and 30 factors for the other datasets. These parameters gave the best results overall (see Supp Figure 8).

### Seurat v4

We compare Mowgli to Seurat v4^15^ which uses Weighted Nearest Neighbors to integrate multi-omics data. We use the R interface Seurat with default parameters.

### Integrative NMF

We implemented a baseline NMF-based integration method by concatenating the features from the different omics and solving the optimization problem with positivity constraints:

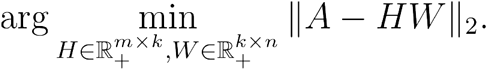

We implemented this approach using the TorchNMF package. As with MOFA+, we chose the number of factors that gave the best results overall (see Supp Figure 9).

Note that this is almost equivalent to intNMF^76^ with *θ* = 1, which minimizes instead 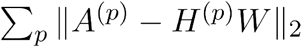. However, on the considered datasets, the intNMF package was too slow to be able to include it in the benchmark.

### Data generation

#### Mixed in RNA

We simulate a dataset where one modality confuses two populations, while the other can separate them. To do so we replace the RNA profiles of all HCT cells with RNA profiles of random HeLa cells. ATAC profiles are left untouched.

#### Mixed in both

We simulate a dataset where the two modalities each confuse two cell populations, but separate two others. This makes the two omics complementary. To do so we replace the RNA profiles of all HCT cells with RNA profiles of random HeLa cells. Then, we replace the ATAC profiles of all K562 cells with ATAC profiles of random HeLa cells.

#### Sparse

We simulate high dropout noise by randomly replacing 50%, 70% ot 90% of the values with zeros. Since the data is already sparse, the final sparsity is 82%, 90% and 96%.

#### Rare population

We simulate the presence of a rare population by keeping only 10 randomly chosen HeLa cells.

### Data preprocessing

All preprocessing was performed using the Scanpy^44^ and Muon^45^ Python packages.

#### RNA preprocessing

Quality control filtering of cells was performed on the proportion of mitochondrial gene expression, the number of expressed genes, and the total number of counts (using Muon’s **filter_obs**). Quality control filtering of genes was performed on the number of cells expressing the gene (using Muon’s **filter_var**). Cells were normalized to sum to 10,000 (using Scanpy’s **normalize_total**), then log-transformed (using Scanpy’s **log1p**). The top 2,500 most variable genes (1,500 for the *Liu* dataset) were selected for downstream analysis (using Scanpy’s **highly_variable_genes** with flavor=‘seurat’).

#### ATAC preprocessing

Quality control filtering of cells was performed on the number of open peaks and the total number of counts (using Muon’s **filter_obs**). Quality control filtering of peaks was performed on the number of cells where the peak is open (using Muon’s **filter_var**). In *Liu, TEA*, and *10X PBMC*, cells were normalized to sum to 10,000 (using Scanpy’s **normalize_total**), then log-transformed (using Scanpy’s **log1p**). In *OP Multiome*, cells were normalized using TF-IDF (using Muon’s **tfidf**) to follow the preprocessing chosen by its authors. The most variable peaks were selected for downstream analysis (using Scanpy’s **highly_variable_genes** with flavor=‘seurat’). Due to differences in the data’s distribution across datasets, we chose to keep 1,500 peaks in *Liu*, 5,000 peaks in PBMC, and 15,000 peaks in *OP Multiome* and *TEA*.

#### ADT preprocessing

Since the number of proteins is small and the data is less noisy than RNA or ATAC, no quality control or feature selection was performed. The data was normalized by Center Log Ratio (using Muon’s **clr**).

### Data analysis

#### Gene Set Enrichment Analysis (GSEA)

The gProfiler API^77^ was used through Scanpy’s **enrich**. Custom sources *GO:CC, GO:MF, GO:BP, Azimuth*, and *ImmuneSigDB* were retrieved from the Enrichr website^78^. Gene sets enriched with adjusted p-values under 0.05 (with Bonferroni correction) were selected for further analysis. To make genes comparable, we normalized rows of the matrix *H*^(*rna*)^ to 1. The 150 top genes for every factor were then used as an unordered input to gProfiler.

#### Motif Enrichment Analysis

Signac^79^ was used to perform Motif Enrichment Analysis, using the JASPAR2022 motif database^80^. To make peaks comparable, we normalized rows of the matrix *H*^(*atac*)^ to 1. The 100 top peaks for every factor were used as input to Signac’s FindMotifs. The union of the top peaks across factors constitutes the background.

#### Visualization

To visualize the latent representation of cells in MOFA+, integrative NMF, and Mowgli’s models, we computed kNN graphs (k = 20) with the euclidean distance between the cells’ low-dimensional embeddings (using Scanpy’s **neighbors**). We used these graphs to compute 2-D UMAP^35^ projections (using Scanpy’s **umap**). For Seurat v4, 2-D UMAP projections based on WNN graphs were performed using Seurat v4’s function RunUMAP.

#### Clustering

For Mowgli, integrative NMF, and MOFA+, we clustered datasets using the Leiden algorithm^36^ with varying resolutions (using Scanpy’s **leiden**). Similarly to UMAP visualization, the inputs of the Leiden algorithm were the previously computed kNN graphs. For Seurat v4, Leiden clustering was performed using Seurat v4’s function FindClusters.

### Evaluation metrics

#### Silhouette score

For each sample, the silhouette width is defined as 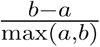 where *a* is the mean distance of the sample to other samples of the same cluster and the *b* is the mean distance of the samples to samples from the nearest cluster. The silhouette score is the mean of silhouette widths across samples. The silhouette score varies between −1 and 1. We used Scikit-learn’s implementation **silhouette_score**^81^.

#### kNN purity score

The kNN purity score measures the average proportion of a sample’s nearest neighbors that share the sample’s cluster annotation. It thus varies between 0 and 1.

#### Adjusted Rand Index

The Rand Index defines the similarity between a ground truth annotation and an experimental clustering. The ARI is then defined as

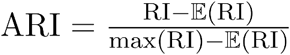

and varies between −1 and 1, with 0 representing a random clustering. We used Scikit-learn’s implementation **adjusted_rand_score**.

### Specificity

MOFA+ was applied with 15 factors, which are enough to represent the data (see Supp Figure 8). Mowgli was applied with 50 factors. In both datasets, a coarse Leiden clustering was applied (using Scanpy’s **leiden** with resolution 0.2). In both datasets, each cluster was assigned a cell type based on the expression of the canonical gene and protein markers (see Supp Figure 1). To confirm this annotation, Azimuth was run on the RNA signal of the dataset (using the Azimuth web tool and the PBMC reference). The agreement of the three independent annotations is confirmed in a Sankey diagram (see Supp Figure 2). Dendritic cells are absent in our manual annotations because of the coarseness of the clustering. Likewise, the ADT signal (see Supp Figure 5) informs us that there is a CD4-CD8-T cell population missed in all three annotations. For each factor in Mowgli and MOFA+ and each cell type, we computed (i) the proportion of cells within that cell type with an absolute weight greater than 10^−3^ (ii) *a*, the mean weight for cells within that cell type (iii) *b*, the mean weight for cells outside of that cell type. For each cell type, we then defined a specificity score for factor *i*:

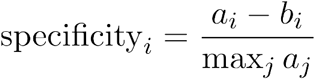

The specificity score is thus bounded by 1. See Figure 4C for a visualization of this information.

### Biological interpretation

We added stars in front of biologically interesting elements in Figure 5C. The first resource we used was the Human Protein Atlas, from which we programmatically retrieved information about the top proteins and genes. We starred them if they were marked as specific to NK cells, Naive B cells, Memory B cells, or Memory CD8 T cells respectively. In addition, some genes or proteins were starred manually; we discuss those in the Results and refer to the relevant literature.

We starred gene sets if they matched the considered cell types, e.g. *MHC II protein complex* for B cells. To reduce the noise in the Immune gene sets, we only considered gene sets opposing subtypes of the broad cell type considered, e.g. *NAIVE_VS_MEMORY_BCELL_UP*.

We starred the TFs if they target one of the top 20 genes. For this, we retrieved TF-gene links from the Regulatory Circuits database^82^ and considered the *Natural Killer cells, CD19+ B cells*, and *CD8+ T cells* networks.

## Availability of data and materials

### Package

The Python package for Mowgli is hosted at https://github.com/cantinilab/mowgli/ and can be installed easily by running `pip install mowgli`.

### Reproducibility

Code to reproduce the experiments and figures is available at https://github.com/cantinilab/mowgli_reproducibility/.

### Regulatory Circuits

At the time of writing, the Regulatory Circuits website http://ww1.regulatorycircuits.org/ is down. We recovered the data from the mirror http://www2.unil.ch/cbg/regulatorycircuits/FANTOM5_individual_networks.tar.

### PBMC

We retrieve a 10X Genomics Multiome (RNA + ATAC) dataset with 9,320 PBMCs. Data is available at https://www.10xgenomics.com/resources/datasets/pbmc-from-a-healthy-donor-granulocytes-removed-through-cell-sorting-10-k-1-standard-2-0-0.

### Liu

We retrieve a scCAT-seq (RNA + ATAC) dataset from Liu et al.^8^ with 206 cells from three cancer cell lines (HCT116, HeLa-S3, K562). Data is available in the Supplementary Materials of the original publication.

### OP-Multiome

We retrieve a Multiome (RNA + ATAC) dataset from the Open Problems challenge^50^ and select only the first batch, which contains 6,137 BMMCs. The GEO accession number is GSE194122 and the data is available at https://www.ncbi.nlm.nih.gov/geo/query/acc.cgi?acc=GSE194122.

### OP-CITE

We retrieve a CITE-seq (RNA + ADT) dataset from the Open Problems challenge^50^ and select only the first batch, which contains 4,249 BMMCs. The GEO accession number is GSE194122 and the data is available at https://www.ncbi.nlm.nih.gov/geo/query/acc.cgi?acc=GSE194122.

### BM-CITE

We retrieve a CITE-seq (RNA + ADT) dataset from Stuart et al.^25^ with 29,803 BMMCs. The GEO accession number is GSE128639 and the data is available at https://www.ncbi.nlm.nih.gov/geo/query/acc.cgi?acc=GSE128639.

### PBMC TEA-seq

We retrieve a recent TEA-seq (RNA + ATAC + ADT) dataset from Swanson et al.^7^ with 7,084 PBMCs. The GEO accession number is GSE158013 and the data is available at https://www.ncbi.nlm.nih.gov/geo/query/acc.cgi?acc=GSE158013.

## Acknowledgements

We would like to thank Frank Augé and Tommaso Andreani for the insightful scientific discussions on this project in the context of the Sanofi iTech Awards.

## Author contribution

G-J.H., G.P. and L.C. designed and planned the study. G-J.H.and L.C. wrote the paper. G.P. revised the manuscript. G-J.H. developed the tool and performed all the analyses. IM.D. participated in the data analysis.

## Ethics declaration

### Ethics approval and consent to participate

Not applicable

## References

1. Rajewsky, N. et al. LifeTime and improving European healthcare through cell-based interceptive medicine. Nature 587, 377–386 (2020).

2. Potter, S. S. Single-cell RNA sequencing for the study of development, physiology and disease. Nat. Rev. Nephrol. 14, 479–492 (2018).

3. Papalexi, E. & Satija, R. Single-cell RNA sequencing to explore immune cell heterogeneity. Nat. Rev. Immunol. 18, 35–45 (2018).

4. Lee, J., Hyeon, D. Y. & Hwang, D. Single-cell multiomics: technologies and data analysis methods. Exp. Mol. Med. 52, 1428–1442 (2020).

5. Stoeckius, M. et al. Simultaneous epitope and transcriptome measurement in single cells. Nat. Methods 14, 865–868 (2017).

6. Clark, S. J. et al. scNMT-seq enables joint profiling of chromatin accessibility DNA methylation and transcription in single cells. Nat. Commun. 9, 781 (2018).

7. Swanson, E. et al. Simultaneous trimodal single-cell measurement of transcripts, epitopes, and chromatin accessibility using TEA-seq. eLife 10, e63632 (2021).

8. Liu, L. et al. Deconvolution of single-cell multi-omics layers reveals regulatory heterogeneity. Nat. Commun. 10, 470 (2019).

9. Angermueller, C. et al. Parallel single-cell sequencing links transcriptional and epigenetic heterogeneity. Nat. Methods 13, 229–232 (2016).

10. Cao, J. et al. Joint profiling of chromatin accessibility and gene expression in thousands of single cells. Science 361, 1380–1385 (2018).

11. Chen, S., Lake, B. B. & Zhang, K. High-throughput sequencing of the transcriptome and chromatin accessibility in the same cell. Nat. Biotechnol. 37, 1452–1457 (2019).

12. Mimitou, E. P. et al. Multiplexed detection of proteins, transcriptomes, clonotypes and CRISPR perturbations in single cells. Nat. Methods 16, 409–412 (2019).

13. Miao, Z., Humphreys, B. D., McMahon, A. P. & Kim, J. Multi-omics integration in the age of million single-cell data. Nat. Rev. Nephrol. 17, 710–724 (2021).

14. Argelaguet, R. et al. MOFA+: a statistical framework for comprehensive integration of multi-modal single-cell data. Genome Biol. 21, 111 (2020).

15. Hao, Y. et al. Integrated analysis of multimodal single-cell data. Cell 184, 3573–3587.e29 (2021).

16. Gayoso, A. et al. Joint probabilistic modeling of single-cell multi-omic data with totalVI. Nat. Methods 18, 272–282 (2021).

17. Ashuach, T., Gabitto, M. I., Jordan, M. I. & Yosef, N. MultiVI: deep generative model for the integration of multi-modal data. http://biorxiv.org/lookup/doi/10.1101/2021.08.20.457057 (2021) doi:10.1101/2021.08.20.457057.

18. Zuo, C. & Chen, L. Deep-joint-learning analysis model of single cell transcriptome and open chromatin accessibility data. Brief. Bioinform. 22, bbaa287 (2021).

19. Duren, Z. et al. Regulatory analysis of single cell multiome gene expression and chromatin accessibility data with scREG. Genome Biol. 23, 114 (2022).

20. Singh, R., Hie, B. L., Narayan, A. & Berger, B. Schema: metric learning enables interpretable synthesis of heterogeneous single-cell modalities. Genome Biol. 22, 131 (2021).

21. Wang, X. et al. BREM-SC: a bayesian random effects mixture model for joint clustering single cell multi-omics data. Nucleic Acids Res. 48, 5814–5824 (2020).

22. Jin, S., Zhang, L. & Nie, Q. scAI: an unsupervised approach for the integrative analysis of parallel single-cell transcriptomic and epigenomic profiles. Genome Biol. 21, 25 (2020).

23. Kim, H. J., Lin, Y., Geddes, T. A., Yang, J. Y. H. & Yang, P. CiteFuse enables multi-modal analysis of CITE-seq data. Bioinformatics 36, 4137–4143 (2020).

24. Welch, J. D. et al. Single-Cell Multi-omic Integration Compares and Contrasts Features of Brain Cell Identity. Cell 177, 1873–1887.e17 (2019).

25. Stuart, T. et al. Comprehensive Integration of Single-Cell Data. Cell 177, 1888–1902.e21 (2019).

26. Stanojevic, S., Li, Y., Ristivojevic, A. & Garmire, L. X. Computational Methods for Single-cell Multi-omics Integration and Alignment. Genomics Proteomics Bioinformatics (2022) doi:10.1016/j.gpb.2022.11.013.

27. Ainsworth, S., Foti, N., Lee, A. K. & Fox, E. Interpretable VAEs for nonlinear group factor analysis. Preprint at http://arxiv.org/abs/1802.06765 (2018).

28. Svensson, V., Gayoso, A., Yosef, N. & Pachter, L. Interpretable factor models of singlecell RNA-seq via variational autoencoders. Bioinformatics 36, 3418–3421 (2020).

29. Lee, D. D. & Seung, H. S. Learning the parts of objects by non-negative matrix factorization. Nature 401, 788–791 (1999).

30. Monge, G. Memoire sur la theorie des deblais et des remblais. Mem Math Phys Acad R. Sci 666–704 (1781).

31. Huizing, G.-J., Peyré, G. & Cantini, L. Optimal transport improves cell–cell similarity inference in single-cell omics data. Bioinformatics 38, 2169–2177 (2022).

32. Stein-O’Brien, G. L. et al. Enter the Matrix: Factorization Uncovers Knowledge from Omics. Trends Genet. 34, 790–805 (2018).

33. Cantini, L. et al. Benchmarking joint multi-omics dimensionality reduction approaches for the study of cancer. Nat. Commun. 12, 1–12 (2021).

34. Maaten, L. van der & Hinton, G. Visualizing Data using t-SNE. J. Mach. Learn. Res. 9, 2579–2605 (2008).

35. McInnes, L., Healy, J., Saul, N. & Großberger, L. UMAP: Uniform Manifold Approximation and Projection. J. Open Source Softw. 3, 861 (2018).

36. Traag, V. A., Waltman, L. & van Eck, N. J. From Louvain to Leiden: guaranteeing well-connected communities. Sci. Rep. 9, 5233 (2019).

37. Blondel, V. D., Guillaume, J.-L., Lambiotte, R. & Lefebvre, E. Fast unfolding of communities in large networks. J. Stat. Mech. Theory Exp. 2008, P10008 (2008).

38. Mootha, V. K. et al. PGC-1α-responsive genes involved in oxidative phosphorylation are coordinately downregulated in human diabetes. Nat. Genet. 34, 267–273 (2003).

39. Korhonen, J. H., Palin, K., Taipale, J. & Ukkonen, E. Fast motif matching revisited: high-order PWMs, SNPs and indels. Bioinformatics 33, 514–521 (2017).

40. Rolet, A., Cuturi, M. & Peyré, G. Fast dictionary learning with a smoothed Wasserstein loss. in Artificial Intelligence and Statistics 630–638 (PMLR, 2016).

41. Qian, W., Hong, B., Cai, D., He, X. & Li, X. Non-Negative Matrix Factorization with Sinkhorn Distance. in IJCAI 1960–1966 (2016).

42. Schmitz, M. A. et al. Wasserstein dictionary learning: Optimal transport-based unsupervised nonlinear dictionary learning. SIAM J. Imaging Sci. 11, 643–678 (2018).

43. Zhang, S. Y. A unified framework for non-negative matrix and tensor factorisations with a smoothed Wasserstein loss. in 2021 IEEE/CVF International Conference on Computer Vision Workshops (ICCVW) 4178–4186 (IEEE, 2021). doi:10.1109/ICCVW54120.2021.00466.

44. Wolf, F. A., Angerer, P. & Theis, F. J. SCANPY: large-scale single-cell gene expression data analysis. Genome Biol. 19, 15 (2018).

45. Bredikhin, D., Kats, I. & Stegle, O. MUON: multimodal omics analysis framework. Genome Biol. 23, 42 (2022).

46. Lähnemann, D. et al. Eleven grand challenges in single-cell data science. Genome Biol. 21, 31 (2020).

47. Qiu, P. Embracing the dropouts in single-cell RNA-seq analysis. Nat. Commun. 11, 1169 (2020).

48. Lance, C. et al. Multimodal single cell data integration challenge: Results and lessons learned. in Proceedings of the NeurIPS 2021 Competitions and Demonstrations Track 162–176 (PMLR, 2022).

49. Luecken, M. D. et al. Benchmarking atlas-level data integration in single-cell genomics. Nat. Methods 19, 41–50 (2022).

50. Luecken, M. et al. A sandbox for prediction and integration of DNA, RNA, and proteins in single cells. in Proceedings of the Neural Information Processing Systems Track on Datasets and Benchmarks (eds. Vanschoren, J. & Yeung, S.) vol. 1 (2021).

51. Lanier, L. L. NKG2D receptor and its ligands in host defense. Cancer Immunol. Res. 3, 575–582 (2015).

52. Boles, K. S., Barchet, W., Diacovo, T., Cella, M. & Colonna, M. The tumor suppressor TSLC1/NECL-2 triggers NK-cell and CD8+ T-cell responses through the cell-surface receptor CRTAM. Blood 106, 779–786 (2005).

53. Prince, H. E., York, J. & Jensen, E. R. Phenotypic comparison of the three populations of human lymphocytes defined by CD45RO and CD45RA expression. Cell. Immunol. 145, 254–262 (1992).

54. Shah, K., Al-Haidari, A., Sun, J. & Kazi, J. U. T cell receptor (TCR) signaling in health and disease. Signal Transduct. Target. Ther. 6, 1–26 (2021).

55. Intlekofer, A. M. et al. Effector and memory CD8+ T cell fate coupled by T-bet and eomesodermin. Nat. Immunol. 6, 1236–1244 (2005).

56. Pirron, U., Schlunck, T., Prinz, J. C. & Rieber, E. P. IgE-dependent antigen focusing by human B lymphocytes is mediated by the low-affinity receptor for IgE. Eur. J. Immunol. 20, 1547–1551 (1990).

57. Bartee, E., Mansouri, M., Hovey Nerenberg, B. T., Gouveia, K. & Früh, K. Downregulation of Major Histocompatibility Complex Class I by Human Ubiquitin Ligases Related to Viral Immune Evasion Proteins. J. Virol. 78, 1109–1120 (2004).

58. Glass, D. R. et al. An Integrated Multi-omic Single-Cell Atlas of Human B Cell Identity. Immunity 53, 217–232.e5 (2020).

59. Lukin, K., Fields, S., Hartley, J. & Hagman, J. Early B cell factor: Regulator of B lineage specification and commitment. Semin. Immunol. 20, 221–227 (2008).

60. Kaileh, M. & Sen, R. NF-κB function in B lymphocytes. Immunol. Rev. 246, 254–271 (2012).

61. Schroeder, H. W. & Cavacini, L. Structure and function of immunoglobulins. J. Allergy Clin. Immunol. 125, S41–S52 (2010).

62. Ody, C. et al. Junctional adhesion molecule C (JAM-C) distinguishes CD27+ germinal center B lymphocytes from non-germinal center cells and constitutes a new diagnostic tool for B-cell malignancies. Leukemia 21, 1285–1293 (2007).

63. Weber, C., Fraemohs, L. & Dejana, E. The role of junctional adhesion molecules in vascular inflammation. Nat. Rev. Immunol. 7, 467–477 (2007).

64. Doñate, C. et al. Homing of Human B Cells to Lymphoid Organs and B-Cell Lymphoma Engraftment Are Controlled by Cell Adhesion Molecule JAM-C. Cancer Res. 73, 640–651 (2013).

65. Laidlaw, B. J. & Cyster, J. G. Transcriptional regulation of memory B cell differentiation. Nat. Rev. Immunol. 21, 209–220 (2021).

66. Vivier, E., Tomasello, E., Baratin, M., Walzer, T. & Ugolini, S. Functions of natural killer cells. Nat. Immunol. 9, 503–510 (2008).

67. Roda-Navarro, P. et al. Human KLRF1, a novel member of the killer cell lectin-like receptor gene family: molecular characterization, genomic structure, physical mapping to the NK gene complex and expression analysis. Eur. J. Immunol. 30, 568–576 (2000).

68. Su, B., Bochan, M. R., Hanna, W. L., Froelich, C. J. & Brahmi, Z. Human granzyme B is essential for DNA fragmentation of susceptible target cells. Eur. J. Immunol. 24, 2073–2080 (1994).

69. Guo, H., Cruz-Munoz, M.-E., Wu, N., Robbins, M. & Veillette, A. Immune Cell Inhibition by SLAMF7 Is Mediated by a Mechanism Requiring Src Kinases, CD45, and SHIP-1 That Is Defective in Multiple Myeloma Cells. Mol. Cell. Biol. 35, 41–51 (2015).

70. Zhang, J. et al. Sequential actions of EOMES and T-BET promote stepwise maturation of natural killer cells. Nat. Commun. 12, 5446 (2021).

71. Ponti, C. et al. Role of CREB transcription factor in c-fos activation in natural killer cells. Eur. J. Immunol. 32, 3358–3365 (2002).

72. Bernard, K. et al. Engagement of natural cytotoxicity programs regulates AP-1 expression in the NKL human NK cell line. J. Immunol. Baltim. Md 1950 162, 4062–4068 (1999).

73. Kantorovich, L. On the transfer of masses (in Russian). in Doklady Akademii Nauk vol. 37 227–229 (1942).

74. Cuturi, M. Sinkhorn Distances: Lightspeed Computation of Optimal Transport. in Advances in Neural Information Processing Systems (eds. Burges, C. J., Bottou, L., Welling, M., Ghahramani, Z. & Weinberger, K. Q.) vol. 26 (Curran Associates, Inc., 2013).

75. Paszke, A. et al. PyTorch: An Imperative Style, High-Performance Deep Learning Library. in Advances in Neural Information Processing Systems vol. 32 (Curran Associates, Inc., 2019).

76. Chalise, P. & Fridley, B. L. Integrative clustering of multi-level ‘omic data based on non-negative matrix factorization algorithm. PLOS ONE 12, e0176278 (2017).

77. Raudvere, U. et al. g:Profiler: a web server for functional enrichment analysis and conversions of gene lists (2019 update). Nucleic Acids Res. 47, W191–W198 (2019).

78. Chen, E. Y. et al. Enrichr: interactive and collaborative HTML5 gene list enrichment analysis tool. BMC Bioinformatics 14, 128 (2013).

79. Stuart, T., Srivastava, A., Madad, S., Lareau, C. A. & Satija, R. Single-cell chromatin state analysis with Signac. Nat. Methods 18, 1333–1341 (2021).

80. Castro-Mondragon, J. A. et al. JASPAR 2022: the 9th release of the open-access database of transcription factor binding profiles. Nucleic Acids Res. 50, D165–D173 (2022).

81. Pedregosa, F. et al. Scikit-learn: Machine Learning in Python. J. Mach. Learn. Res. 12, 2825–2830 (2011).

82. Marbach, D. et al. Tissue-specific regulatory circuits reveal variable modular perturbations across complex diseases. Nat. Methods 13, 366–370 (2016).

83. PBMC from a Healthy Donor - Granulocytes Removed Through Cell Sorting (10k). 10x Genomics https://www.10xgenomics.com/resources/datasets/pbmc-from-a-healthy-donor-granulocytes-removed-through-cell-sorting-10-k-1-standard-2-0-0.

