## Supplementary Information for "Paired single-cell multi-omics data integration with Mowgli"

### Supplementary material

| Dataset name | Technology | Modalities | Organism | Tissue | Number of cells | Number of annotations | Reference |
|---|---|---|---|---|---|---|---|
| Liu | scCAT-seq | RNA, ATAC | Human | Cell lines | 206 | 3 | Liu et al.[8] |
| 10X PBMC | 10X Mutliome | RNA, ATAC | Human | PBMC | 9320 | 14 | 10X Genomics[83] |
| OP Multiome | 10X Multiome | RNA, ATAC | Human | Bone Marrow | 6137 | 21 | Open Problems[50] |
| OP CITE | CITE-seq | RNA, ADT | Human | Bone Marrow | 4249 | 35 | Open Problems[50] |
| BM CITE | CITE-seq | RNA, ADT | Human | Bone Marrow | 29803 | 27 | Stuart et al.[25] |
| TEA | TEA-seq | RNA, ATAC, ADT | Human | PBMC | 7084 | no annotations | Swanson et al.[7] |

Supplementary Table 1. List of the datasets used in this paper and their characteristics.

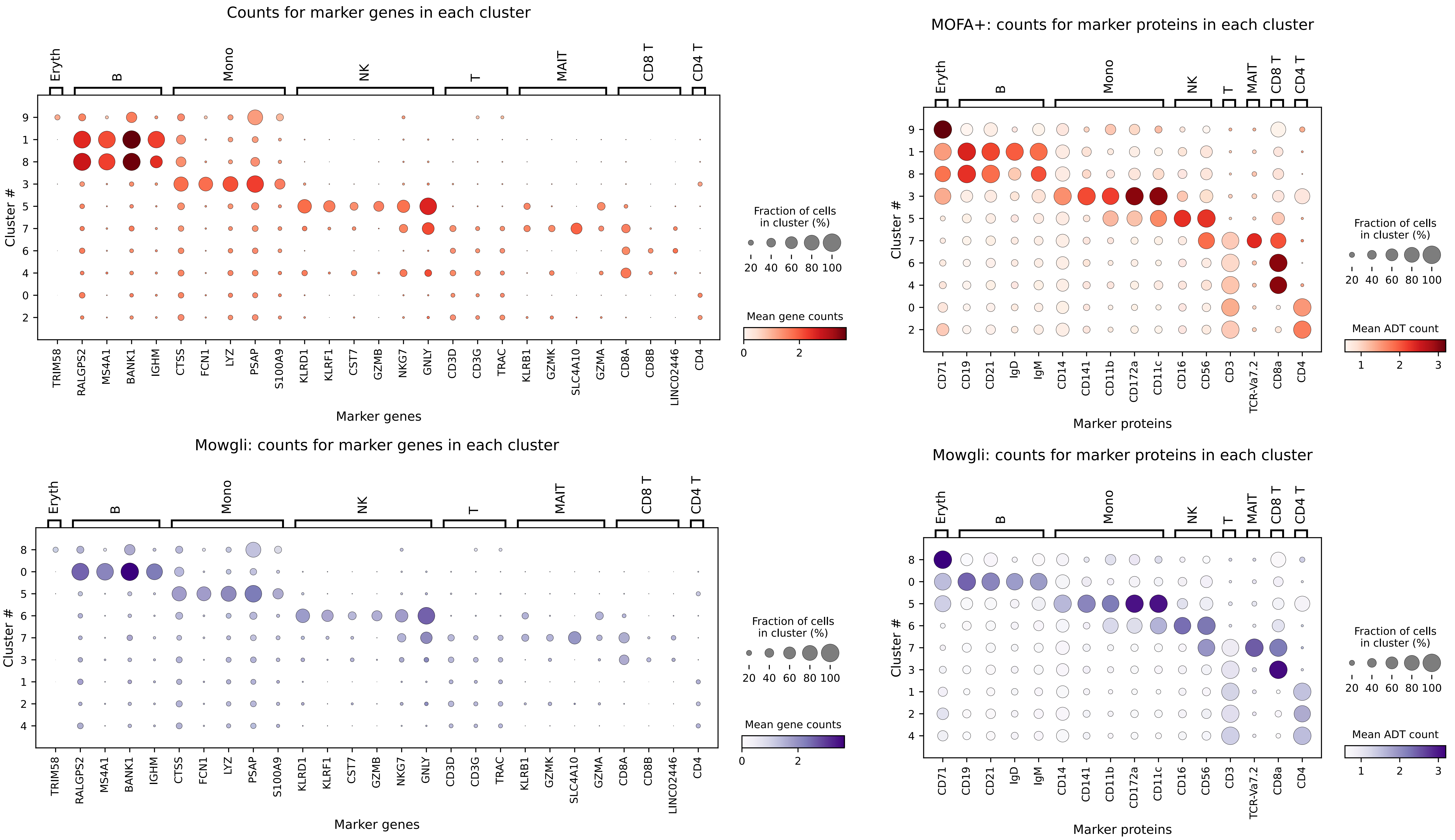


Supplementary Figure 1. Expression of gene and protein markers in the coarse Leiden clusters in the TEA-seq PBMC dataset, for Mowgli and MOFA+.

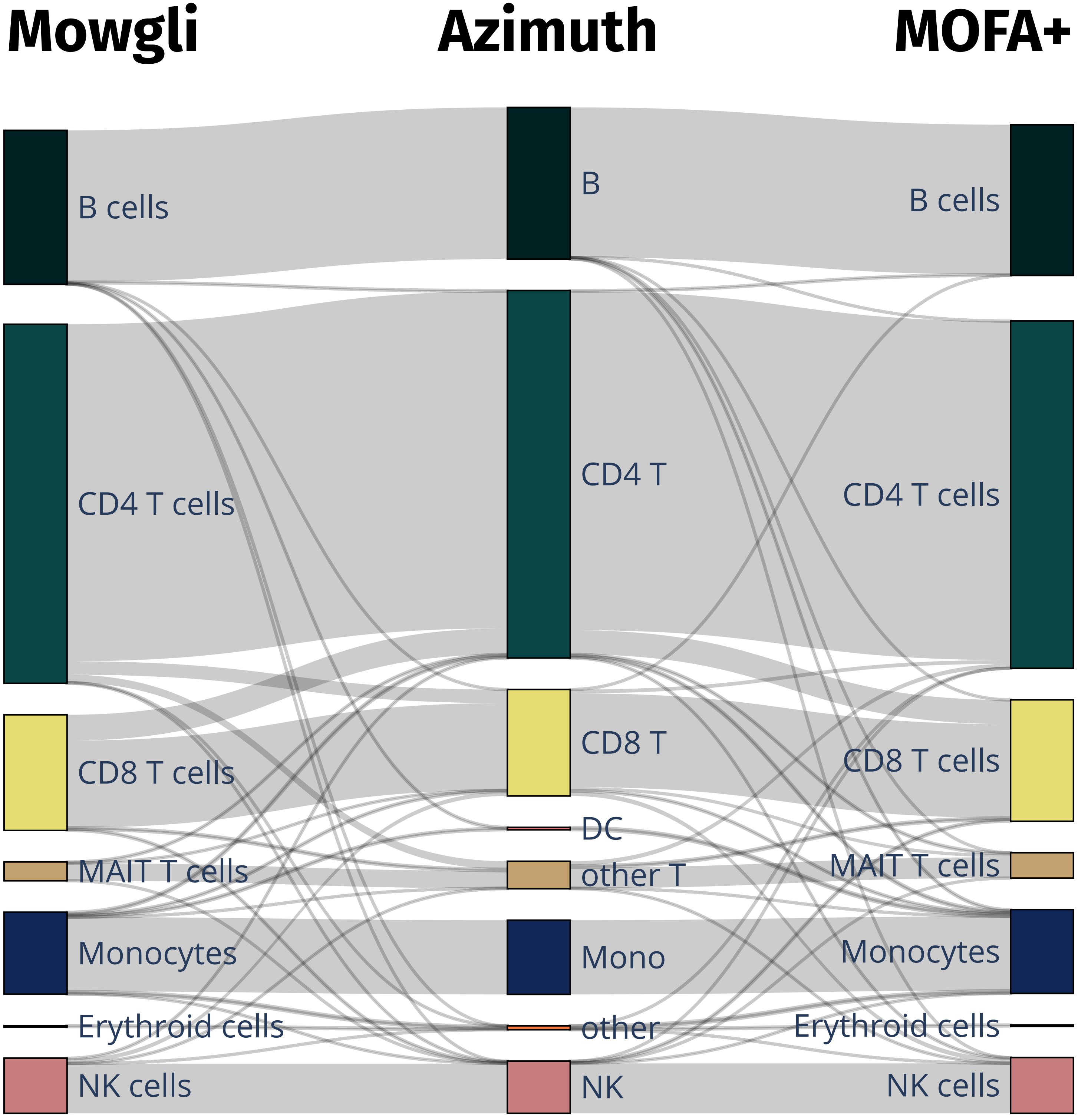


Supplementary Figure 2. Sankey diagram showing the agreement between Mowgli's annotation (left), Azimuth's annotation (middle) and MOFA+'s annotation (right).

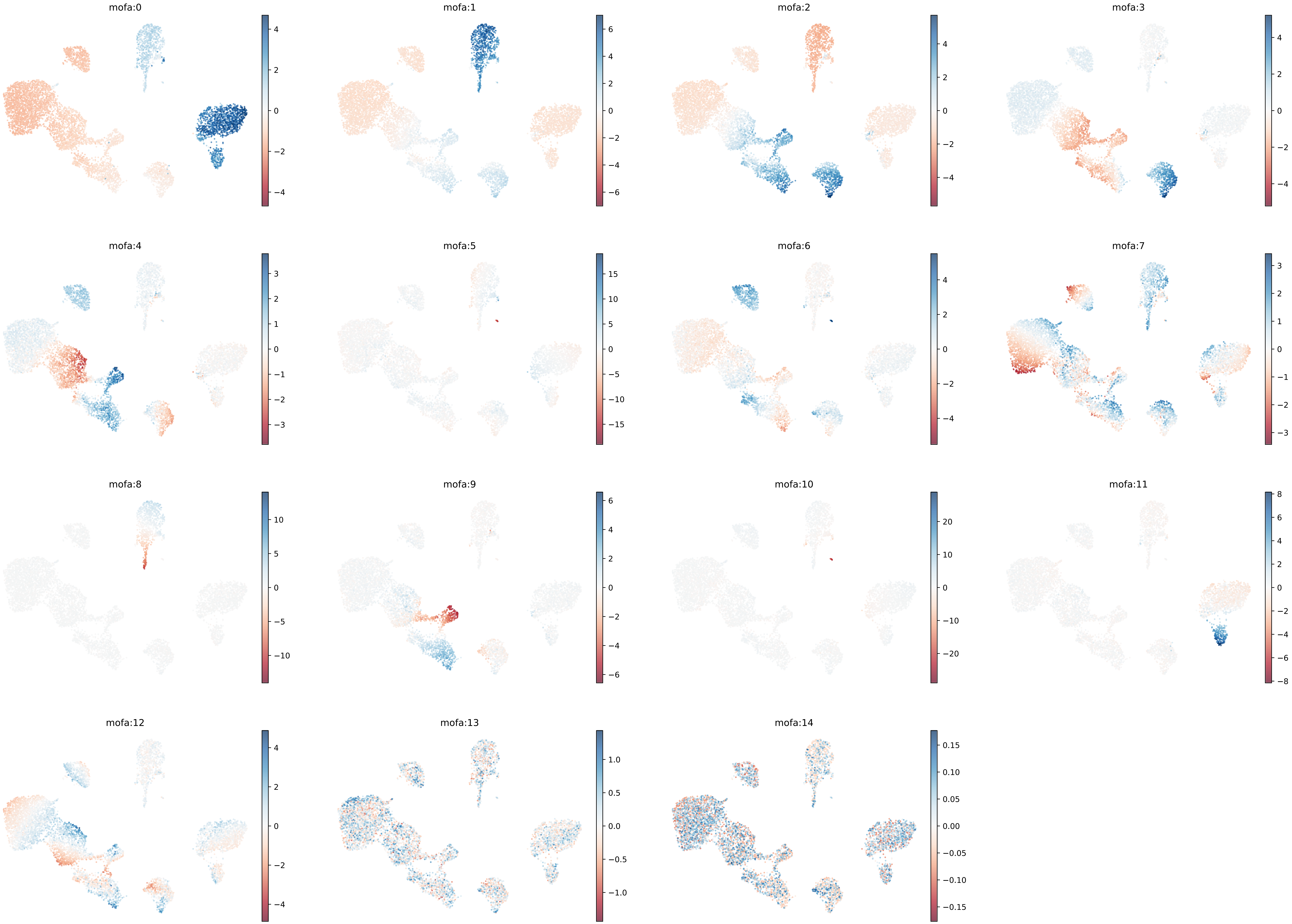


Supplementary Figure 3. UMAP plots with MOFA+'s factors, each cell colored by weight.

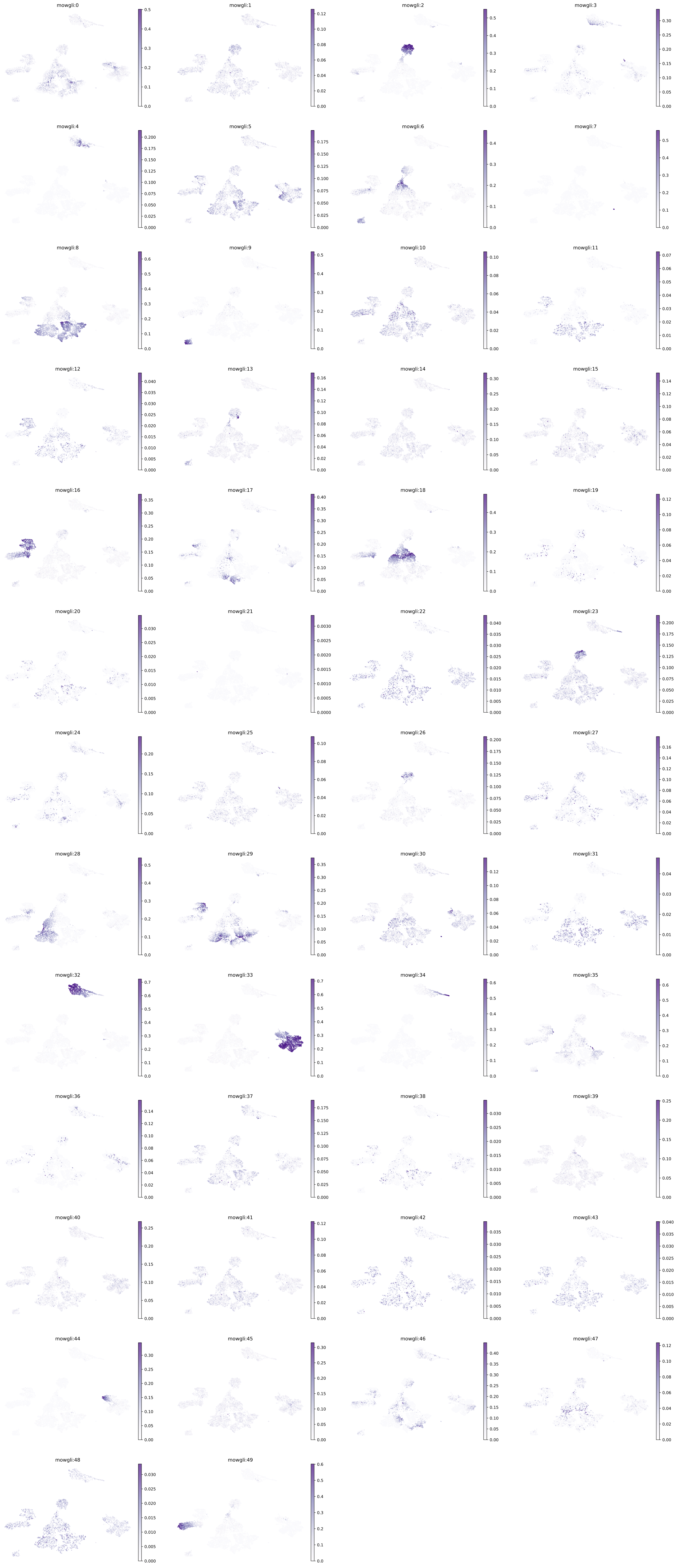


Supplementary Figure 4. UMAP plots with Mowgli’s factors, each cell colored by weight.

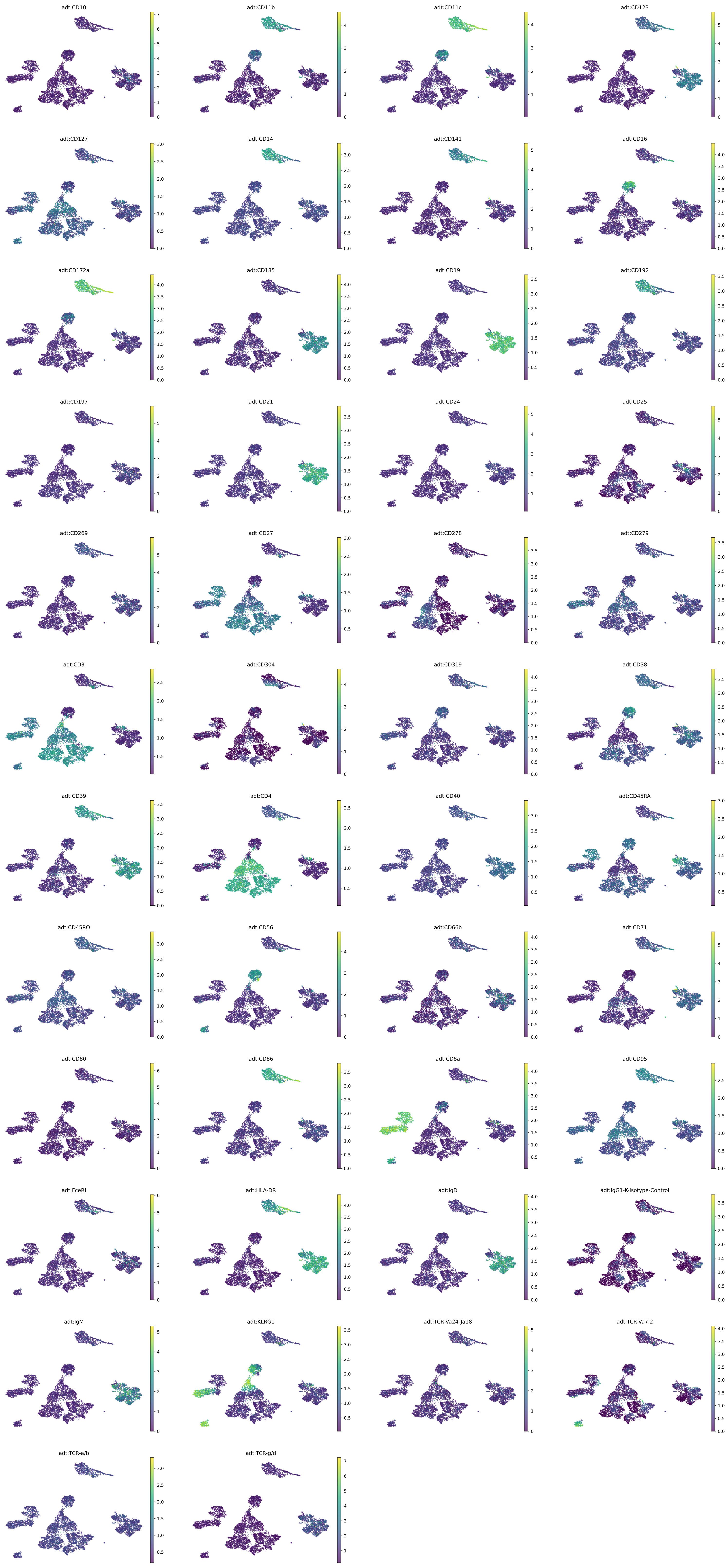


Supplementary Figure 5. UMAP plots of Mowgli cell embeddings, colored by protein counts.

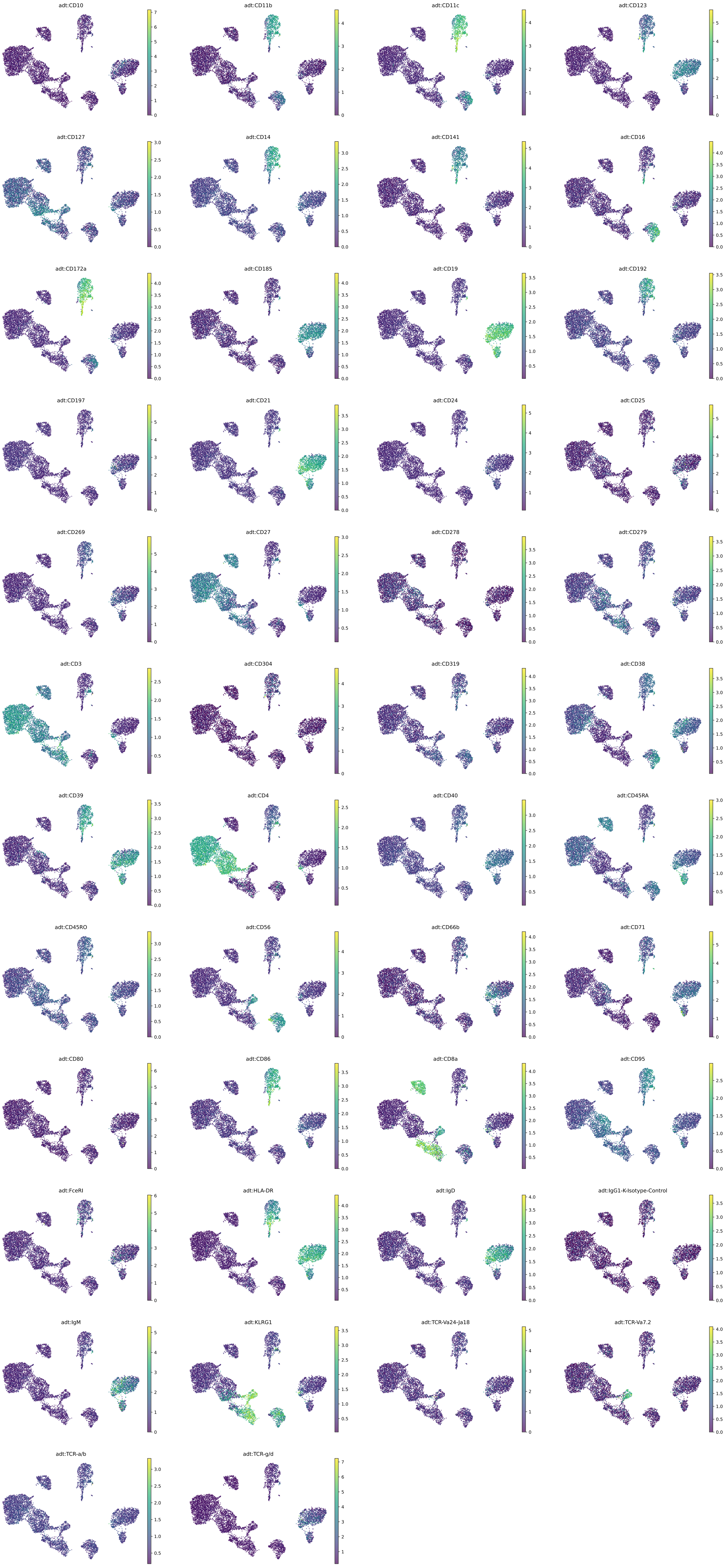


Supplementary Figure 6. UMAP plots of MOFA+ cell embeddings, colored by protein counts.

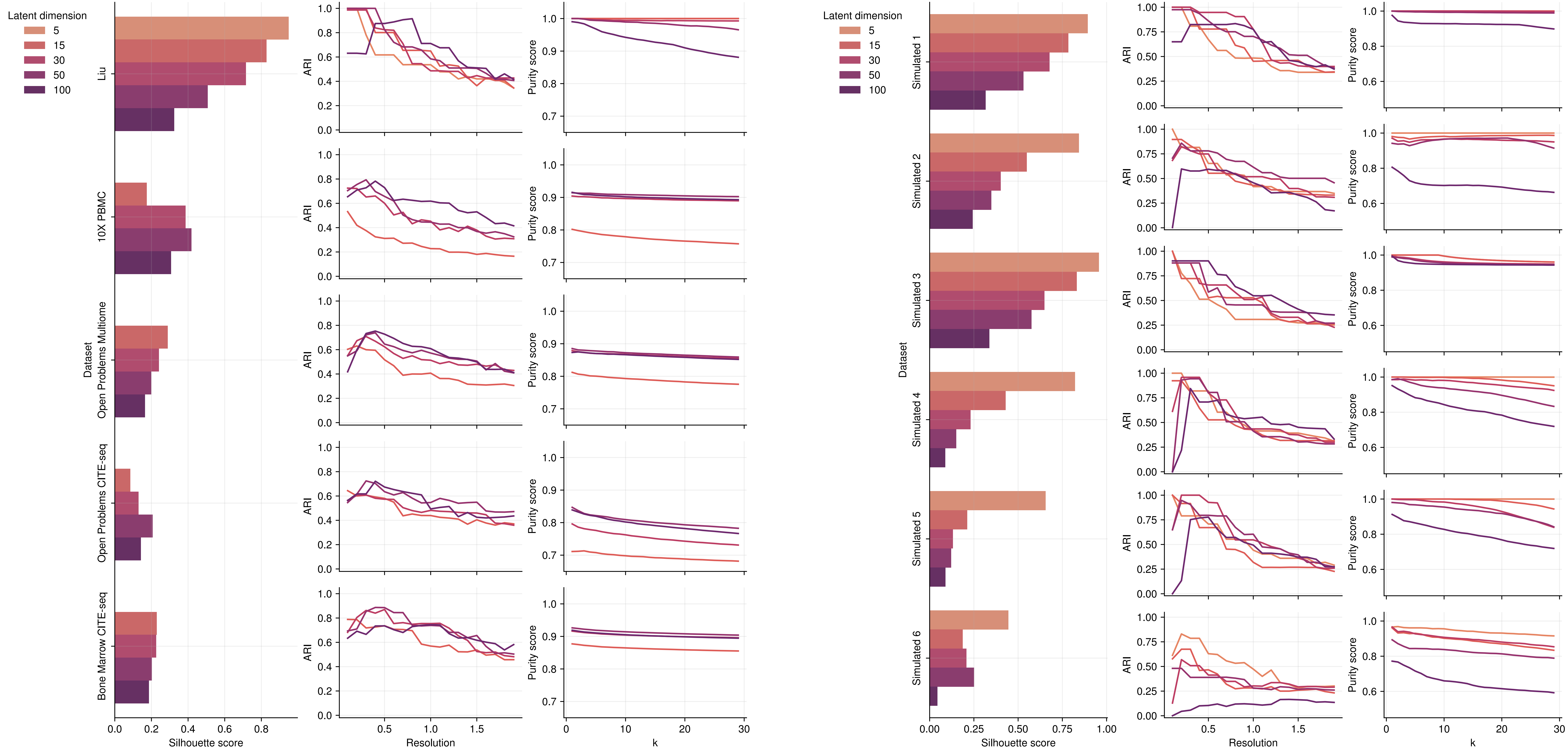


Supplementary figure 7. Mowgli's clustering and embedding quality, evaluated across latent dimensions.

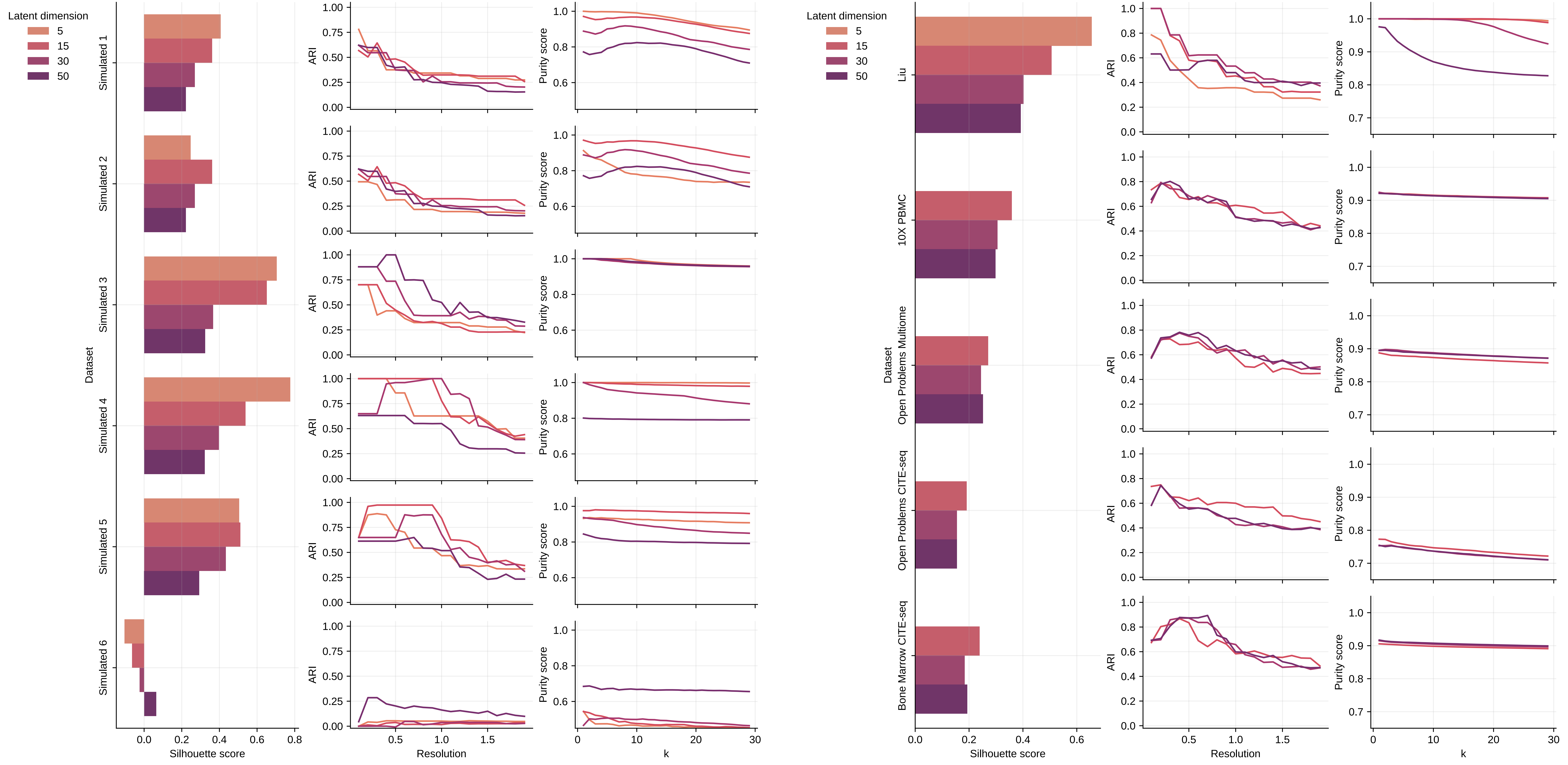


Supplementary figure 8. MOFA+’s clustering and embedding quality, evaluated across latent dimensions.

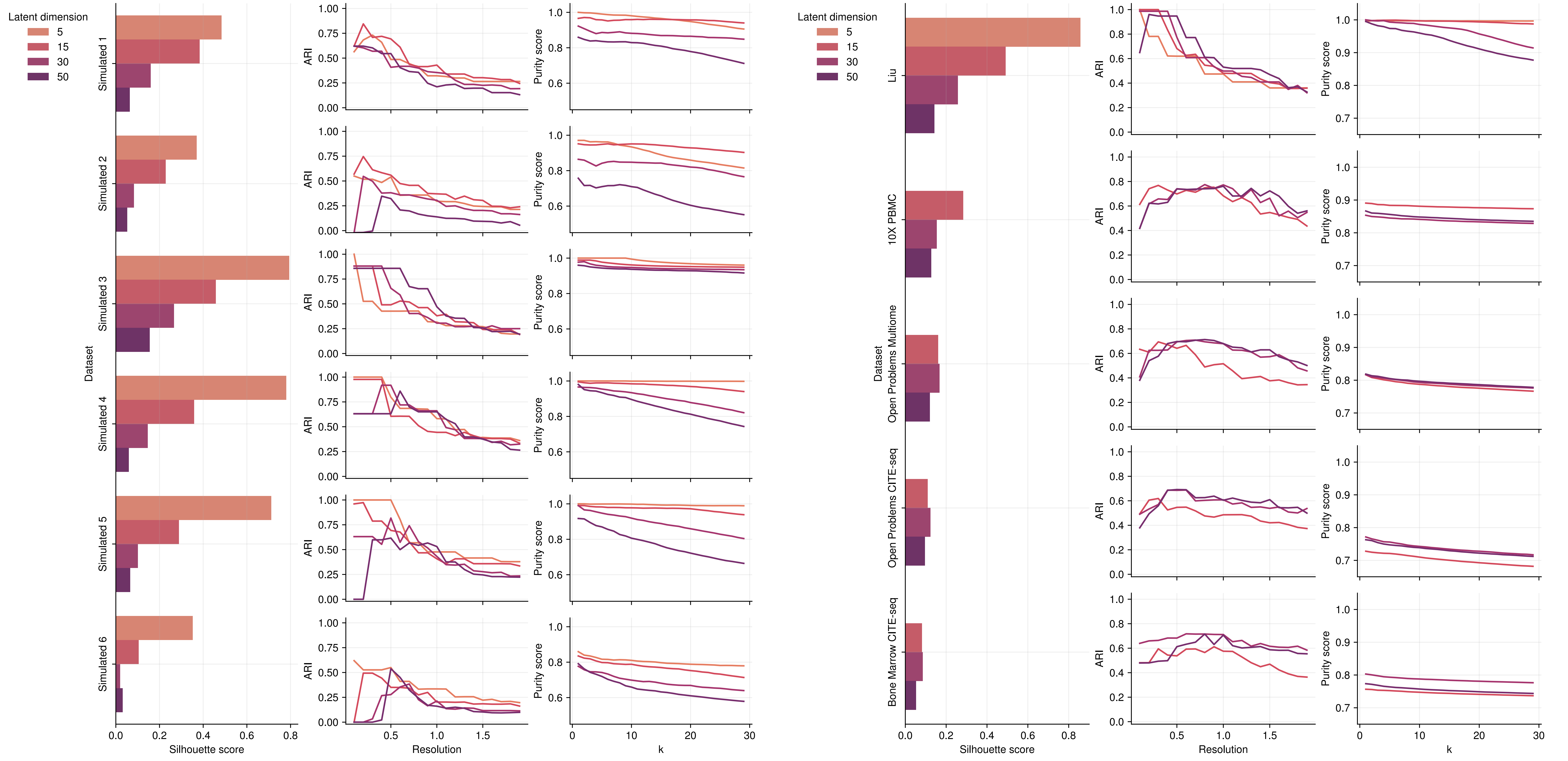


Supplementary figure 9. Integrative NMF's clustering and embedding quality, evaluated across latent dimensions.
